# Impairment of the Hif-1α regulatory pathway in Foxn1-deficient (Foxn1^−/−^) mice affects the skin wound healing process

**DOI:** 10.1101/2020.08.04.237388

**Authors:** Sylwia Machcinska, Marta Kopcewicz, Joanna Bukowska, Katarzyna Walendzik, Barbara Gawronska-Kozak

## Abstract

Hypoxia and hypoxia-regulated factors [e. g., hypoxia-inducible factor-1α (Hif-1α), factor inhibiting Hif-1α (Fih-1), thioredoxin-1 (Trx-1), aryl hydrocarbon receptor nuclear translocator 2 (Arnt-2)] have essential roles in skin wound healing. Using Foxn1^−/−^ mice that can heal skin injuries in a unique scarless manner, we investigated the interaction between Foxn1 and hypoxia-regulated factors. The Foxn1^−/−^ mice displayed impairments in the regulation of Hif-1α, Trx-1 and Fih-1 but not Arnt-2 during the healing process. An analysis of wounded skin showed that the skin of the Foxn1^−/−^ mice healed in a scarless manner, displaying rapid re-epithelialization and an increase in *transforming growth factor β* (*Tgfβ-3)* and *collagen III* expression. An *in vitro* analysis revealed that Foxn1 overexpression in keratinocytes isolated from the skin of the Foxn1^−/−^ mice led to reduced *Hif-1α* expression in normoxic but not hypoxic cultures and inhibited *Fih-1* expression exclusively under hypoxic conditions. These data indicate that in the skin, Foxn1 affects hypoxia-regulated factors that control the wound healing process and suggest that under normoxic conditions, Foxn1 is a limiting factor for Hif-1α.

## INTRODUCTION

Skin, the largest organ of the body, serves as a protective barrier against infections, heat loss, dehydration, and mechanical insults and can sense external (temperature, touch) and internal (neurons, hormones) signals ^1, 2^. Interruption of skin integrity by injury or surgical intervention can lead to severe damage and abnormal homeostasis ^3^. The integrity of the wounded skin is restored through a reparative healing process, which results in scar formation. Although scarring is beneficial for the reestablishment of the protective skin function, it often leads to impairment of skin performance as a sensing organ, and under some circumstances (e.g., burn scars), it may cause major disabilities.

Skin wound healing is initiated by blood clot formation that restores homeostasis and provides a fibrin matrix, which becomes the scaffold for cells infiltrating the wounded area ^4^. The clot and surrounding wound tissues release numerous mediators: cytokines and growth factors such as platelet-derived growth factor (PDGF), transforming growth factor-1 (TGFß-1), fibroblast growth factor-7 (FGF-7) and epidermal growth factor (EGF) that attract macrophages, neutrophils and fibroblasts, initiating the inflammatory phase of healing ^5^. During the proliferative phase, keratinocytes proliferate and migrate over the provisional matrix to cover the wound in the process of re-epithelialization, endothelial cells support angiogenesis (neovascularization) within the wound bed, and fibroblasts/myofibroblasts produce and secrete collagen, proteoglycans and glycosaminoglycans, which are the main components of the extracellular matrix (ECM) ^6-8^. The remodelling phase that begins 2–3 weeks after injury is controlled by matrix metalloproteinases (MMPs) and their tissue inhibitors (TIMPs), which mediate the gradual replacement of collagen III by collagen I. Then, most of the cells in the wound bed (macrophages, endothelial cells and myofibroblasts) undergo programmed cell death, and acellular scar is formed ^3, 7^.

The process of regenerative (scar-free/scarless) skin wound healing, which perfectly restores the architecture and function of the wounded skin, is an uncommon phenomenon among mammals. For a long time, the skin of mammalian foetuses was believed to be the only example of regenerative cutaneous healing ^7, 9^. However, recent data have shown that the skin of adult Foxn1^−/−^ (nude) and Acomys mice is also capable of scarless healing similar to that observed in mammalian foetuses ^10, 11^. The common features of regenerative skin wound healing are rapid re-epithelialization, reduced inflammation and lack of scarring ^9-13^. Moreover, regenerative wounded skin has high levels of collagen III, tenascin C, transforming growth factor β-3 (TGFβ-3) and hyaluronic acid (HA) expression ^10, 11, 13, 14^. Furthermore, during regenerative healing, MMP-9 and MMP-13, metalloproteinases involved in the skin healing process, show a unique, bimodal pattern of expression that is upregulated in the early and late phases of wound healing ^13^. In adult mammals, scarless wound healing has been shown to occur in mice that have a spontaneous, inactivating mutation in the Foxn1 gene ^10, 13, 15^. Holes punched in the external ears of Foxn1^−/−^ mice regrow, and skin injuries heal in a regenerative manner similar to that of mammalian foetuses during the first two trimesters of gestation ^10, 13, 15, 17^. A subsequent study revealed that Foxn1 is a regulator of the reparative (scar-forming) skin wound healing process, participating in re-epithelialization and scar formation due to its activity during epithelial-mesenchymal transition (EMT) ^28, 29^. The expression of the transcription factor Foxn1 has been detected in two organs with multilayered epithelial structures: the thymus and the skin. In the thymus, Foxn1 is involved in T cell development ^18, 19^. In the skin, where Foxn1 is expressed in the epidermis and in hair follicles, it participates in hair follicle development, promotes keratinocyte differentiation and participates in the pigmentation process ^20-23^. Loss-of-function mutations in the *Foxn1* gene has pleiotropic effects, similar in humans and mice and result in a lack of hair and the absence of the thymus and T cells ^24-27^.

Although comprehensive studies have identified several factors and pathways participating in the skin wound healing process, the molecular regulation that allows the body to redirect scar-forming processes to scar-free processes or nonhealing wounds to scar formation remains unclear.

One of the major factors that may guide reparative vs regenerative skin wound healing is hypoxia. Oxygen (O_2_) is essential for aerobic organisms to produce energy via mitochondria and to perform other biological processes and is one of the most dangerous factors to the cell ^30, 31^. However, acute hypoxia in injured tissues, including skin wounds, is a crucial environmental cue that triggers the wound healing process. Restoration of cellular homeostasis in the hypoxic environment is orchestrated by the major player hypoxia-inducible factor-1α (Hif-1α). In tissues, under normal oxygen conditions (normoxia), Hif-1α is rapidly degraded (post-translational process) through a reaction catalysed by HIF prolyl-4-hydroxylases (PHD) and factor inhibiting Hif-1 (Fih-1) in an O_2_–dependent manner ^32, 33^. Under hypoxic conditions, Hif-1α is protected from degradation due to depletion of PHDs and Fih-1 via proteasomal degradation and enhancement of the Hif-1α protein levels by thioredoxin-1 (Trx-1) ^23, 34, 35^. Then, Hif-1α translocates to the nucleus, undergoes dimerization with Hif-1β and activates the transcription of downstream genes, e.g., vascular endothelial growth factor (VEGF), platelet-derived growth factor B-chain (PDGF-B) and heat shock protein (HSP-90) ^32, 36^. Hif-1α, as a master regulator of oxygen homeostasis, contributes to all stages of wound healing: cell migration and proliferation, growth factor release and matrix synthesis.

Hif-1α deficiency leads to chronic hypoxia, which contributes to the formation of nonhealing ulcers ^3^. In contrast, Hif-1α overexpression has been associated with fibrotic disease, and elevated Hif-1α protein levels were detected in keloids and scleroderma tissues ^3^. These data indicate that the hypoxic environment and the expression levels and modulation of hypoxia-related genes (e.g., Hif-1α) direct the skin wound healing response ^32, 37-40^. Furthermore, mammalian foetal skin wounds showed rapid and scar-free healing (regenerative) in a physiologically hypoxic environment ^41^.

Recently, using next-generation high-throughput DNA sequencing, we performed detailed analyses of differentially expressed genes in the skins of Foxn1^−/−^ and Foxn1^+/+^ (wild type) mice and found significant differences in the expression of genes associated with hypoxia ^23^. Furthermore, the *in vitro* experiments showed that under low-oxygen conditions, Foxn1 expression is induced in primary cultures of keratinocytes ^23^.

In the present study, we tested the hypothesis that the interaction between Foxn1 and hypoxia-regulated factors during cutaneous wound healing drives the physiological decision between regeneration (scarless healing) or repair (scarring).

## MATERIALS AND METHODS

### Animals

The studies were performed on mature, 4–6-month-old Foxn1^−/−^ (nude mice; CBy.Cg-Foxn1<nu>/cmdb) and genetically matched controls Foxn1^+/+^ (Balb/c/cmdb) that were housed in individually ventilated cages (IVC) in a temperature- and humidity-controlled room (22°C and 55%, respectively) with a 12 h light/12 h dark cycle at the Center of Experimental Medicine (CEM), Medical University of Bialystok, Poland. The experimental animal procedures were approved by the Ethics Committee of the University of Warmia and Mazury (Olsztyn, Poland), No. 68/2018. The study was carried out in accordance with EU Directive 2010/63/EU of the European Parliament and of the Council on the protection of animals used for scientific purposes (OJEU, 2010. Official Journal of the European Union. Directive 2010/63/EU of the European Parliament and of the Council on the protection of animals used for scientific purposes. OJEU. [cited 2010 Oct 20]; Series L 276:33-79.).

### Experiment I (Supplementary Figure 1)

The day before the experiment, Foxn1^+/+^ mice were anaesthetized with isoflurane, and the dorsal area was shaved and rinsed with an alcohol swab. The next day, Foxn1^+/+^ and Foxn1^−/−^ mice were anaesthetized by isoflurane, and sterile 4-mm diameter biopsy punch (Miltex, GmbH, Rietheim-Weilheim, Germany) was used to create 4 excisional wounds on the backs of the mice. Excised skin samples (4 mm in diameter) were collected, immediately frozen in liquid nitrogen and stored at −80°C until analysis (day 0, uninjured control). After wounding, the mice were placed in warmed cages until recovery. Skin samples were collected post-mortem with an 8 mm biopsy punch at days 1, 3, 5, 7, 14, 21 and 36 post-injury (n=6 animals per time point). Excised, wounded skin tissues were frozen in liquid nitrogen for protein and RNA isolation and hydroxyproline content determination. Samples for histological examination were fixed in 10% formalin (FA) (Sigma-Aldrich) or 4% paraformaldehyde (PFA)(Sigma-Aldrich).

**Figure 1.**
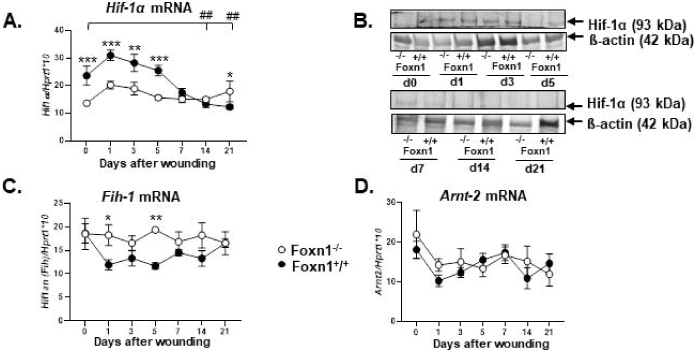
Skin injury evokes weaker *Hif-1α* and *Fih-1* responses in Foxn1^−/−^ than Foxn1^+/+^ mice. *Hif-1α* **(A)**, *Fih-1* (**C**) and *Arnt-2* **(D)** mRNA expression determined by qRT-PCR analysis of uninjured (day 0) and injured (days 1, 3, 5, 7, 14, 21) skin tissues collected from the Foxn1^**−/−**^ and Foxn1^**+/+**^ mice (n=6 skin samples per group). **(B)** Representative Western blot analysis of Hif-1α and β-actin (reference protein) in the skin of the Foxn1^**−/−**^ and Foxn1^**+/+**^ mice. Values are the lsmean ± SE; asterisks indicate significant differences between the Foxn1^**−/−**^ and Foxn1^**+/+**^ mice during the skin wound healing process (*p < 0.05;** p < 0.01; ***p < 0.001), # indicates significant differences between day 0 vs 14 and day 0 vs 21 (^##^ p <0.01) for the Foxn1^**+/+**^ mice.

### Experiment II (Supplementary Figure 2)

We used the bromodeoxyuridine (BrdU) labelling approach to identify and estimate cell proliferation in the wounded skin tissues. The day before the experiment, shaved mice were anaesthetized and given four full-thickness 4 mm diameter wounds. At days 1, 3, 5 and 7 after wounding (n=4 for day 1 and n=2 per days 3, 4 and 7), the mice were anaesthetized and injected (intraperitoneal) with BrdU [dissolved in 4 mg BrdU (Sigma-Aldrich) in 300 µl of saline/per mouse]. The wounded skin tissues were collected 2 or 4 h post-mortem after BrdU injection. Cell isolates were analysed by flow cytometry to identify and estimate the BrdU-labelled cell proliferation rate in the wounded skin tissues. The most effective BrdU labelling was obtained 2 h after BrdU injection. For the main experiment, the Foxn1^−/−^ animals (CBy.Cg-Foxn1<nu>/cmdb; n=28) and genetically matched Foxn1^+/+^ controls (Balb/c/cmdb; n=28) were used. The mice were injured as described in experiment I. BrdU injection was given 2 h before animal sacrifice at days 1, 2, 3, 5 and 7 post-wounding. Skin tissues were collected at day 0 (n=3 as a control) and days 1, 2, 3, 5 and 7 post-wounding (n=5 per time point). Skin samples were collected post-mortem with an 8 mm biopsy punch: one 8 mm punch from each animal was collected for histological analysis, and 3 punches were collected for cell isolation followed by flow cytometric analysis.

**Figure 2.**
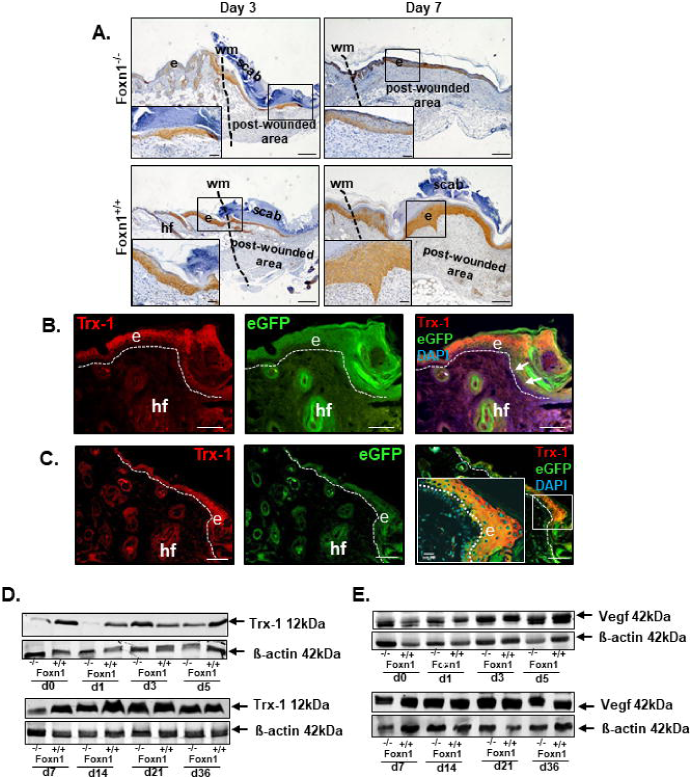
Strong Trx-1 protein accumulation in the skin of the Foxn1^+/+^ mice. **(A)** Immunohistochemical detection of Trx-1 localization in the skin of the Foxn1^**−/−**^ and Foxn1^**+/+**^ mice collected at days 3 and 7 post-injury. **(B-C)** Colocalization of Trx-1 and Foxn1 was performed on skin samples from the transgenic Foxn1::eGFP mice at day 3. Western blot analysis of Trx-1 **(D)** and Vegf **(E)** in the skin of the Foxn1^**−/−**^ and Foxn1^**+/+**^ mice. e – epidermis, wm – wound margin, hf – hair follicles. Scale bar **(A)** 200 µm, (insets 50 µm); **(B)** 100 µm and **(C)** 100 µm (confocal images; inset 20 µm).

### Analysis of wound closure and re-epithelialization

The wound healing process was monitored macroscopically. Photographs were taken at days 1, 3, 5, 7, 14, 21 and 36 after wounding. ImageJ software [National Institutes of Health (NIH) Image] was used to measure wound closure. Wound size [mm^2^] was calculated based on the area of an ellipse = radius of the length × radius of the width × π (n=42), as described previously ^28^. The re-epithelialization process was assessed microscopically on cytokeratin 16-stained histological slides. Measurements were performed with Cell Sens Dimension Software (Olympus Soft Imaging Solutions GmbH) and calculated according to the following formula: (length of the extending epidermal tongues)/(length of the wound) x 100%, as described previously ^28^.

### Histology

Formalin-fixed skin samples were processed, embedded in paraffin and sectioned at 3 µm. Slides for collagen detection were stained with Masson’s trichrome using a standard protocol (Trichrome Stain Kit, Abcam). Immunohistochemical staining for the presence of Ki67 (1:50, Cat# ab16667, Abcam), CD68 (1:200,Cat# 125212, Abcam), cytokeratin 16 (1:300, Cat# b7609, LsBio), Trx-1 (1:200, Cat# 24295, Cell Signaling) and incorporation of BrdU (1:200, Cat# 6326, Abcam) was performed on consecutive skin sections. Antibody binding was detected with the ABC complex (Vectastain ABC kit, Vector Laboratories, Inc.). In the control sections, primary antibodies were substituted with nonspecific immunoglobulin G (IgG). Peroxidase activity was revealed using 3,3′-diaminobenzidine (Sigma-Aldrich) as a substrate. The slides were counterstained with haematoxylin (Sigma-Aldrich). The sections were visualized using an Olympus microscope (BX43), photographed with an Olympus digital camera (XC50) and analysed with Olympus CellSens Software. Immunofluorescence detection and colocalization of Foxn1 and Trx-1 were performed on skin samples from young Foxn1::eGFP transgenic mice at day 3 post-wounding ^42^. The samples were fixed in 4% PFA for 2 h, washed in 0.1 M PB (Sigma-Aldrich) overnight at 4°C and stored in 18% sucrose (Chempur, Poland)/0.1 M PB solution prior to cryosectioning. Immunofluorescence assays were performed with the following primary antibodies: anti-Trx-1 (1:50, Cat# 24295, Cell Signaling) and anti-eGFP (1:400, Cat#6673, Abcam). Alexa Fluor 594(1:200, Cat# A11037, Invitrogen) and Alexa Fluor 488 (1:200, Cat# A 21206, Invitrogen) secondary antibodies were used. Nuclei were counterstained with ProLong Gold Antifade Mountant with DAPI (Invitrogen). The sections were visualized and photographed with an Olympus microscope (BX43) equipped with an Olympus digital camera (XC50) and analysed with CellSens Dimension Software (Olympus Soft Imaging Solutions GmbH). Confocal images were scanned and digitalized using an F10i Laser Scanning Microscope integrated with Fluoview Software (Olympus) with a 10x and 60x objective lens. The sequential scans were acquired with Z spacing of 0.5 μm and 512 x 512 pixel size at room temperature.

### Protein isolation and Western blot analysis

Frozen skin samples collected at days 0, 1, 3, 5, 7, 14, 21 and 36 were powdered in liquid nitrogen using a prechilled mortar and pestle and then homogenized in RIPA buffer containing protease inhibitor cocktail (Sigma-Aldrich), phosphatase inhibitor cocktail (Sigma-Aldrich) and phenylmethanesulfonyl fluoride (PMSF, Sigma-Aldrich), followed by sonication (3 × 5 s) with a sonicator (Vibro-Cell VCX 130 PB). Protein concentration was measured by the infrared (IR)-based protein quantitation method with a Direct Detect Infrared Spectrometer (Merck). Fifty or thirty-five (depending on the protein detection) micrograms of proteins was separated on 9% Tricine gels and transferred to polyvinylidene difluoride membranes (Merck Millipore). The membranes were incubated separately with anti-Mmp-9 (1:1000, Cat# AB19016, Merck Millipore), anti-Vegf (1:200, Cat# 46154, Abcam), anti-Hif 1-α (1:200, Cat# 16066, Abcam), anti-Trx-1 (1:1000, Cat# 24295, Cell Signaling) or anti-β-actin (1:1000, Cat# 8226, Abcam), followed by incubation with fluorescent secondary antibodies against IRDye 800 (1:10000, Cat# 611-132-122, Rockland Immunochemicals, Inc., PA, USA) or Cy5.5 (1:10000, Cat# 610-113-121, Rockland Immunochemicals, Inc., PA, USA) for 1 h. Bands were visualized using the ChemiDoc™ Touch Imaging System (Bio-Rad) and analysed using Image Lab Software (Bio-Rad) according to the manufacturer’s protocol.

### Hydroxyproline and MCP-1 analysis

Eight millimetre diameter frozen skin punches were homogenized in 1 ml of phosphate buffered saline (PBS, Sigma-Aldrich). Five hundred microlitres of homogenized tissue was frozen at −20°C for Monocyte Chemoattractant Protein-1 (MCP-1) analysis, and the remaining 500 µl was stored overnight at 4°C for the hydroxyproline assay. The next day, 0.5 ml aliquots were hydrolysed with 0.25 ml of 6 N HCl for 5 h at 120°C. For construction of a standard curve, hydroxyproline concentrations from 0 to 20 μg/ml were used (Sigma-Aldrich). Twenty microlitres of each experimental sample and standard curve sample was added to a 96-well plate and incubated for 20 min at room temperature with 50 μl of chloramine T solution ^43^. Then, 50 μl of Ehrlich’s solution (2.5 g 4-(dimethyloamino) benzaldehyde, 9.3 ml n-propanol and 3.9 ml 70% perchloric acid) was added, and the plate was incubated for 15 min at 65°C. After the samples were cooled, the plate was read at 550 nm on a microplate reader (Multiskan Sky Microplate Spectrophotometer, Thermo Fisher Scientific™).

For the (MCP-1) detection assay, frozen homogenates were thawed and centrifuged (5 min/5000 x g, 4°C). The supernatant was collected, aliquoted and stored at −80°C until the assay was performed. The assay procedure was performed according to the manufacturer’s protocol (Cusabio Technology LLC., USA)

### Flow cytometric analysis

Flow cytometry was performed on uninjured (n=3 as a control) and injured (n=5) skin samples from the Foxn1^−/−^ and Foxn1^+/+^ mice collected at 1, 2, 3, 5 and 7 days after wounding. The skin tissues were washed in 70% ethanol and PBS (with 1% penicillin and streptomycin), minced and digested in 3.68 mg/ml collagenase (Sigma-Aldrich) for 80 min at 37°C/160 rpm. Then, the cells were filtered through 100 µm and 70 μm cell strainers (Falcon, A Corning Brand, NY, USA). Next, the cells were centrifuged at 1200 rpm for 5 min at room temperature (RT). The pelleted cells were suspended in Dulbecco’s modified Eagle’s medium (DMEM/F-12; Sigma-Aldrich) supplemented with 15% foetal bovine serum (FBS; Life Technologies) and 1% antibiotics (penicillin/streptomycin, Sigma-Aldrich) and counted in a Burker’s chamber. Next, the cells were centrifuged at 1200 rpm for 5 min at RT and suspended in warm sterile PBS. The assay procedure was performed according to the manufacturer’s protocol [BD Pharmingen BrdU Flow Kit, Becton Dickinson Cat# 559619 (FITC)]. For multiple staining, cells were incubated with APC (Lynx Rapid APC antibody conjugation kit, Cat # LNK031APC, Bio-Rad Laboratories, Inc.)-conjugated cytokeratin 6 (CK6; NSJ Bioreagents, Cat # V2168) antibodies, anti-CD45-PE-Cy7 (BD Pharmingen, Cat # 552848), anti-CD44-PE (BD Pharmingen, Cat # 553134) and anti-CD68-PE (BD Pharmingen, Cat # 566387) antibodies. The labelled cells were analysed using a BD LSRFortessa Cell Analyser flow cytometer (Becton Dickinson) and BD FACSDiva v6.2 Software (Becton Dickinson). The data are expressed as the percentage of BrdU-, CK6-, CD68-, and CD44-positive cells per gated cell.

### RNA isolation and real-time PCR

Total RNA was extracted from the skin samples using TRI Reagent (Invitrogen by Thermo Fisher Scientific). The RNA concentration and quality were determined spectrophotometrically using a NanoDrop 1000 (Thermo Fisher Scientific) and agarose gel electrophoresis. cDNA was synthesized from 500 ng of total RNA using a High-Capacity cDNA Reverse Transcription Kit with RNase Inhibitor (Applied Biosystems by Thermo Fisher Scientific). For measurement of the mRNA levels of *Arnt-2*(Cat# Mm00476009_*m1), Fih-1(Cat# Mm01198376_m1), Foxn1(Cat# Mm01298129), Collagen I* (Cat# Mm00483888), *Collagen III*(Cat# Mm01254476_m1*), Mmp-9* (Cat#, Mm00442991_m1*), Timp-1*(Cat# Mm00441818_m1), *Tgfβ-1*(Cat# Mm01178820), *Tgfβ-3*(Cat# Mm00436960) alpha-SMA (Cat# Mm00725412) and *Vegf* (Cat# Mm00437306_m1) Single Tube TaqMan® Gene Expression Assays (Life Technologies Thermo Fisher Scientific) were used. Hypoxanthine phosphoribosyltransferase 1 (*Hprt1*, Cat# Mm01545399_m1) was chosen as the most stable housekeeping gene during cutaneous wound healing after a previously described analysis ^29^. All samples were analysed in duplicate. Each run included a standard curve based on aliquots of pooled skin. RNA amplification was performed using a 7900HT Fast Real-Time PCR System under the following conditions: initial denaturation for 10 min at 95°C, followed by 40 cycles of 15 s at 95°C and 1 min at 60°C. The mRNA expression was normalized to that of the reference gene *Hprt1* and multiplied by 10.

### Enzyme-linked immunosorbent assay (ELISA) of collagen type I

Dermal fibroblasts (DFs) were isolated from skin samples of 4–6-month-old Foxn1^−/−^ (CBy. Cg-Foxn1<nu>/cmdb) and Foxn1^+/+^ (Balb/c/cmdb) mice (n= 4 samples per group) The samples were mechanically processed and digested in collagenase type 1 (3.68 mg/ml; Sigma-Aldrich) for 1 h and 20 min at 37°C in a shaker platform. Isolates were filtered with a cell strainer with a pore diameter of 100 μm and centrifuged for 5 min at 1300 rpm at room temperature. Pellets were resuspended in culture medium DMEM/F12 (Sigma-Aldrich) supplemented with 15% FBS with antibiotics (penicillin/streptomycin; Sigma-Aldrich) and counted in a Countess™ automated cell counter (Invitrogen). DFs were seeded at 3.0×10^5^ cells in 12-well plates in DMEM/F12 with 15% FBS and 1% antibiotics (penicillin/streptomycin). After the cells reached 50–60% confluency, the culture medium was replaced with fresh DMEM/F12 with 2% FBS supplemented with TGFβ1/TGFβ3 (10 ng/ml; PeproTech EC, Ltd., London, UK) or 5mM citric acid (diluent for lyophilized TGFβ; negative control). Conditioned media were collected after 24, 48 and 72 h of stimulation and stored at −20°C for analysis of collagen concentration. All treatments were performed using cells isolated from five separate pools of skin. Concentrations of collagen I in the culture media were determined using the Enzyme-Linked Immunosorbent Assay Kit for Col1a1 (Cloud Clone Corp.) according to the manufacturer’s protocol. For immunofluorescence detection of Alpha-Smooth Muscle Actin (α-SMA) or vimentin, DFs cultured in DMEM/F12 with 2% FBS supplemented with TGFβ1 or TGFβ3 (10 ng/ml; PeproTech EC, Ltd., London, UK) or 5 mM citric acid (diluent for lyophilized TGFNβ; negative control) were fixed for 15 min in methanol (−20°C) and washed 3 times in PBS. DFs were incubated with primary antibodies against α-SMA (1:200, Cat# RB9010-P, Thermo Fisher Scientific) or Vimentin (1:400, Cat# 92547, Abcam) for 1 h at room temperature. Then, the cells were washed 3 times in PBS and incubated for 1 h in the dark with the following secondary antibodies: Alexa Fluor 594 (1:200, Cat# A11037, Invitrogen) or Alexa Fluor 488 (1:200, Cat# A 21206, Invitrogen). Cells were sealed with ProLong1 Gold Antifade Mountant with DAPI (Life Technologies). In the control sections, primary antibodies were substituted with nonspecific IgG. Cells were visualized and photographed with a confocal microscope (Fluoview FV10i, Olympus).

### Keratinocyte-DF coculture experiments

Skin tissues collected from newborn Foxn1^−/−^ (Cby.Cg-Foxn1<nu>/cmdb) and Foxn1^+/+^ (C57BL/6; B6) mice were incubated in dispase (6 U/ml; Life Technologies) overnight at 4°C. The separated epidermis was digested in 0.05% trypsin-EDTA (Life Technologies) for 3 min and filtered through a 70 μm strainer (Falcon, A Corning Brand, NY, USA). Then, keratinocytes were collected by a series of three trypsin digestions (at 37°C) and filtration and were then centrifuged at 300×g for 9 min at room temperature. The pelleted cells were suspended and seeded in DMEM/F-12 (Sigma-Aldrich) supplemented with 10% FBS (Life Technologies), 0.2% Primocin (InvivoGen, France), and 120 μM β-mercaptoethanol (Sigma-Aldrich). The media were exchanged for CnT medium (CELLnTEC, Switzerland) 24 h after seeding. The remaining dermal tissues were digested for 45 min in collagenase type I (220 U/ml; Life Technologies), filtered through a 70 μm strainer and centrifuged. Isolated DFs were seeded in DMEM/F-12 medium containing 15% FBS and antibiotics (penicillin/streptomycin, Sigma-Aldrich). All *in vitro* experiments were performed on primary (p=0) keratinocytes seeded in inserts at densities of 2.0 ×10^6^ cells and DFs seeded in 6-well plates at a density of 3.0 ×10^5^ cells.

Foxn1^−/−^ (Cby.Cg-Foxn1<nu>/cmdb) keratinocytes at 70% confluency were transduced with Ad-Foxn1 or control Ad-GFP adenovirus (a kind gift from Janice L. Brissette; Harvard Medical School, Boston, MA). Adenoviral transductions were performed at a multiplicity of infection of 200 in 400 µl of CnT basal medium (CELLnTEC, Switzerland). After 4 h, 600 µl of CnT basal medium with supplements A, B, and C (CELLnTEC, Switzerland) was added per insert. Then, the Ad-Foxn1- or Ad-GFP-transduced Foxn1^−/−^ keratinocytes were used for coculture experiments with DFs (cultured on the bottoms of 6-well plates) and cultured for 24 h under normoxic (21% O_2_) or hypoxic (1% O_2_) conditions in a humidified incubator at 37°C. Non-transduced Foxn1^+/+^ (B6) keratinocytes (cultured in inserts) and DFs (cultured on the bottoms of 6-well plates) were established as coculture units and cultured for 12 h or 24 h under normoxic (21% O_2_) or hypoxic (1% O2) conditions in a humidified incubator with 1% O_2_ (37°C).

### Statistical analysis

For determination of the impact of group (Foxn1^−/−^ vs Foxn1^+/+^ mice) and time (days 0–36) on skin parameters (*in vivo* experiments) and the impact of treatment (*in vitro* experiments), linear models were used, in which interaction between these covariates was taken into account as well. Based on such models, lsmeans were calculated (least-squared means, means calculated based on the model’s coefficients) and compared. For each model, the adjusted R squared and p-value of the F-test of overall significance are provided as the measure of goodness-of-fit. The significance level was set to 0.05. All calculations were performed in R (ver. 3.5.3) using the following packages: emmeans (ver. 1.4.1) and tidyverse (ver. 1.2.1). Statistical analysis was performed by Biostat, Poland (https://www.biostat.com.pl/index_en.php).

## RESULTS

### Skin wounding in Foxn1-deficient mice triggers a reduced hypoxic response

Our previous study showed that Foxn1 expression in keratinocytes is induced by hypoxia and that Foxn1 activation increases the levels of hypoxia-regulated proteins in primary cultures of keratinocytes ^23^. In the present study, we examined skin wound healing in terms of hypoxia-regulated factors using Foxn1^−/−^ mice (Cby.Cg-Fox1<nu>/cmdb; nude) and genetically matched Foxn1^+/+^ (Balb/c/cmdb) control mice.

First, we examined the expression of hypoxia-related factors in uninjured (day 0) and injured (days 1–21) skin (Figure 1, Supplementary Tables 1–3). The intact skin of the Foxn1^−/−^ mice displayed lower *Hif-1α* mRNA expression (p<0.001, Figure 1A, Supplementary Tables 1–2) and protein (Figure 1B) levels than that of the Foxn1^+/+^ mice. During the wound healing process, the *Hif-1α* mRNA levels were significantly lower in the skin of the Foxn1^−/−^ mice than the Foxn1^+/+^ mice at days 1 (p<0.001), 3 (p<0.01) and 5 (p<0.001) post-wounding (Figure 1A, Supplementary Table 1). Interestingly, at day 21, a decrease in the *Hif-1α* mRNA levels was detected in the skin of the Foxn1^+/+^ mice relative to their Foxn1^−/−^ counterparts (p<0.05; Figure 1A, Supplementary Table 1). The Hif-1α protein was present in the extracts obtained from the wounded skin of the Foxn1^−/−^ (days 1–3) and Foxn1^+/+^ (days 1–5) mice (Figure 1B). The Foxn1^−/−^ and Foxn1^+/+^ mice also showed differences in the expression of *Fih-1* (a factor inhibiting Hif-1), which participates in the hypoxia-mediated response. The Foxn1^−/−^ mice demonstrated higher *Fih-1* transcript levels than the Foxn1^+/+^ mice at days 1 (p<0.05) and 5 post-wounding (p<0.01; Figure 1C, Supplementary Table 3). Aryl hydrocarbon receptor nuclear translocator 2 (*Arnt-2*; *Hif-2β*), the hypoxia-mediated signalling element, showed similar mRNA levels for both groups of mice (Figure 1D).

Next, we analysed thioredoxin-1 (Trx-1), the main free radical scavenger that enhances the protein levels of Hif-1α (Figures 2A-2D), and Vegf, a target of Hif-1α (Figure 2E). Immunohistochemical staining of Trx-1 was detected in the epidermis adjacent to the wound margin, in the newly formed epidermis covering the wound and in hair follicles (Figure 2A). However, in the Foxn1^−/−^ mice, in contrast to the Foxn1^+/+^ mice, Trx-1 was limited to the basal and suprabasal layers of the epidermis in immunohistochemical analysis (Figure 2A). The higher Trx-1 protein levels in the skin of the Foxn1^+/+^ mice were confirmed by Western blot analysis (Figure 2D).

Since both Foxn1 and Trx-1 are expressed in epithelial cells of the interfollicular epidermis and in hair follicles, we performed colocalization studies on histological samples collected from the wounded skin of transgenic Foxn1::eGFP mice ^28^. Double fluorescent imaging revealed that the epidermis adjacent to the wounded area, the thickened wound margin and the leading epithelial tongue showed strong colocalization of eGFP (an indirect indicator of Foxn1 expression) and Trx-1 (Figures 2B, 2C). Confocal imaging revealed that Trx-1 and Foxn1 colocalized in the suprabasal layer and in differentiating keratinocytes. Colocalization of Foxn1 and Trx-1 was not observed in the basal layer of the epidermis (Figure 2C). Western blot analysis performed on tissues collected from uninjured and injured skin showed no differences in the Vegf protein levels between the Foxn1^−/−^ and Foxn1^+/+^ mice (Figure 2E).

### Accelerated re-epithelialization is observed in excisional skin wounds of the Foxn1^−/−^ mice

The macroscopic evaluation of full-thickness excisional skin wounds in the Foxn1^−/−^ and Foxn1^+/+^ mice revealed a faster rate of skin wound closure in the Foxn1^−/−^ mice than the Foxn1^+/+^ mice, with substantial differences in wound size at day 3 (p<0.01; Figures 3A, 3B, Supplementary Tables 4–5). To analyse the progress of re-epithelialization, we performed immunohistochemical detection of cytokeratin 16 (CK16), an indicator of activated keratinocytes (migration/proliferation) (Figure 3C). Samples collected from the Foxn1^−/−^ mice exhibited a faster rate of re-epithelialization than those from the Foxn1^+/+^ animals (Figures 3C, 3D, Supplementary Tables 6–7). The morphometric analysis of the distance between the two leading epithelial tongues allowed us to calculate the percentage of wound area covered by newly formed epithelium, which was significantly higher for the Foxn1^−/−^ mice than the Foxn1^+/+^ mice on day 1 (24.1% ± 14.4 vs 6.3% ± 6.4), day 2 (70.3% ± 37.1 vs 20.9%± 8.7; p <0.01) and day 3 (94.2% ± 10.1 vs 80.9% ± 20.9) after injury (Figures 3C, 3D, Supplementary Tables 6–7). Complete re-epithelialization was observed at day 5 post-injury for both groups of mice (Figures 3C, 3D, Supplementary Tables 6–7). Next, we used the BrdU labelling approach (BrdU incorporation assay) to estimate the cell proliferation rate in the wounded skin tissues (Supplementary Figure 2; Figure 4, Supplementary Tables 8–11). Flow cytometric analysis showed that in the uninjured mice, the percentage of skin cells incorporating BrdU was similar for the Foxn1^−/−^ (2.83% ± 0.75) and Foxn1^+/+^ (2.80% ± 0.75) mice (Supplementary Table 9). In contrast, the analysis of cells isolated from the injured mice injected with BrdU revealed substantial differences in the rate of BrdU incorporation between the Foxn1^−/−^ and Foxn1^+/+^ mice. Generally, the Foxn1^−/−^ mice displayed a higher percentage of BrdU^+^/CD45^−^ (CD45 a marker of haematopoietic lineage) skin cells collected at days 1 (3.14% ± 0.58 vs 1.74% ± 0.58), 2 (4.94% ± 0.58 vs 2.40% ± 0.58; p<0.05) and 3 (2.06% ± 0.58 vs 0.48% ± 0.58) than the Foxn1^+/+^ mice (Figures 4A, 4B, Supplementary Table 9). The rate of BrdU incorporation stabilized at days 5 and 7 and became similar for both groups of mice (Figure 4A, Supplementary Table 9). The phenotypic characterization of the BrdU-incorporating cells showed that 40–60% were CD44 positive (marker of cells with mesenchymal origin), regardless of the mouse strain or the day post-injury (Figure 4C). In contrast, the percentage of cytokeratin 6 (CK6)-positive cells in the population of BrdU-labelled cells was higher in the wounded skin of the Foxn1^+/+^ mice than that of the Foxn1^−/−^ mice at days 2 (p<0.05) and 3 (p<0.01; Figure 4D; Supplementary Table 10). To identify the localization of proliferating cells, we performed immunohistochemical detection of BrdU or Ki67 (Figures 4E-4G). Generally, BrdU (Figure 4E) or Ki67 (Figure 4F, 4G) was detected in the epidermis and hair follicles (Figures 4E-4G). A large cluster of BrdU-positive cells was observed in the basal layer and occasionally in the suprabasal layer of the epidermis (Figure 4E). An increased number of BrdU- or Ki67-positive cells in the skin was identified at days 2 and 3 post-wounding (Figures 4E-4G). Interestingly, the accumulation of BrdU-positive cells was detected in the epidermis adjacent to the injured region but not in the neoepidermis (Figure 4E). These data support recent findings by Aragona et al., which showed the presence of two distinct populations of keratinocytes around the wound: cells proximal to the wound edge that migrate and cells distal to the wound edge that proliferate, supplying a new pool of cells that can in turn migrate into the wound ^44^.

**Figure 3.**
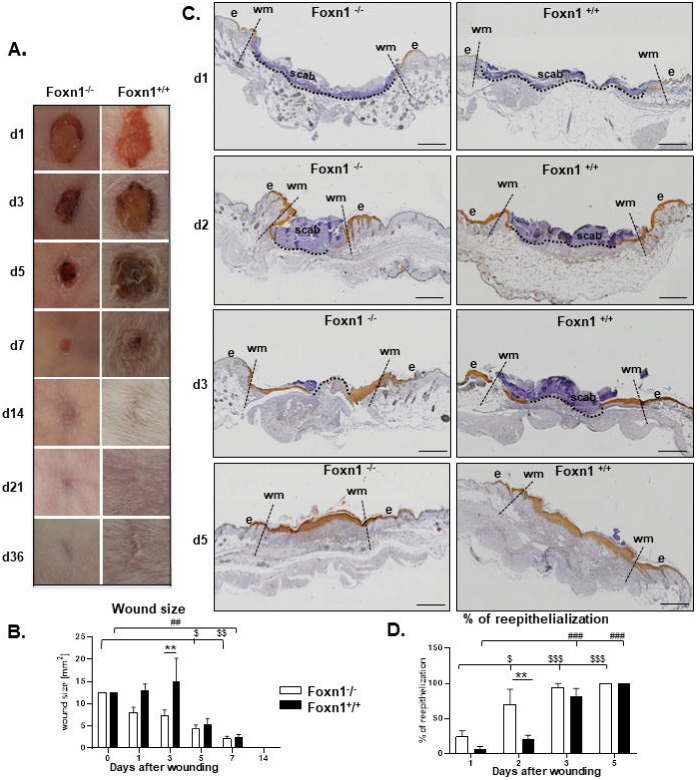
Accelerated skin wound healing in the Foxn1^−/−^ mice. **(A)** Representative macroscopic images of skin wounds at days 1–36 after wounding. **(B)** Morphometrical analysis of the wound closure areas (n=12 measurements per time point). **(C)** Immunohistochemical detection of cytokeratin 16 (CK16) in the wounded skin sections from the Foxn1^−/−^ and Foxn1^+/+^ mice at days 1, 2, 3 and 5. **(D)** Morphometrical analysis of the re-epithelization process in the skin of the Foxn1^−/−^ and Foxn1^+/+^ mice (n=3 mice per group). Wm-wound margin; e-epidermis. **(C)** Scale bar 500 μm. Values are the lsmean ± SE; asterisks indicate significant differences between the Foxn1^−/−^ and Foxn1^+/+^ mice (**p<0.01); # indicates significant differences between day 0 (B) or day 1 (D) and respective days of healing in the Foxn1^+/+^ mice (^##^ p<0.01; ^####^ p<0.001); $ indicates significant differences between day 0 (B) or day 1 (D) and respective days of healing in the Foxn1^**−/−**^ mice (^$^p<0.05; ^$$^p<0.01; ^$$$^p<0.001).

**Figure 4.**
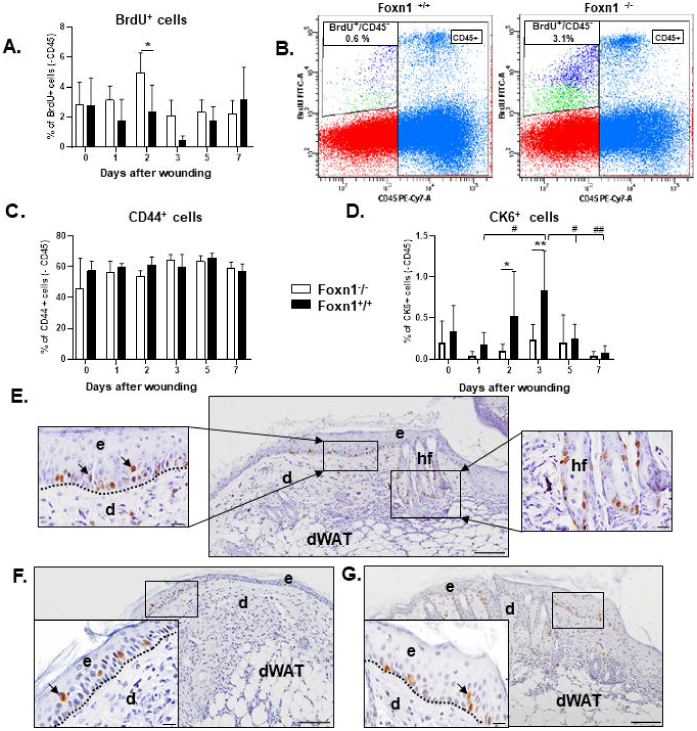
The wounded skin of the Foxn1^−/−^ mice displays a greater proliferative capacity than the skin from the Foxn1^+/+^ mice. Flow cytometry (**A-D)** and immunohistochemical analysis of BrdU **(E)** and Ki67 **(F-G)**-positive cells in the wounded skin collected from the Foxn1^**−/−**^ and Foxn1^**+/+**^ mice injected with BrdU 2 h prior to sacrifice. Percentage of BrdU^**+**^ (CD45^**-**^) cells **(A-B)** in total cell isolates; percentage of CD44^+^ (CD45^**-**^) **(C)** and CK6^+^ (CD45^**-**^) **(D)** cells in the BrdU-positive population (n=3–5 per group). Immunohistochemical detection of BrdU-positive cells in the skin of the Foxn1^**+/+**^ mice on day 3 **(E)** and Ki67-positive cells in the skin of Foxn1^**−/−**^ on day 2 **(F)** and Foxn1^**+/+**^ on day 3 **(G)**. e-epidermis, d-dermis, dWAT-dermal white adipose tissue, hf –hair follicle. Scale bars: 100 µm, insets 20 µm. Values are the lsmean ±SE; asterisks indicate significant differences between the Foxn1^**−/−**^ and Foxn1^**+/+**^ mice. (*p < 0.05;** p < 0.01); # indicates the differences between day 1 vs day 3 and day 3 vs day 5 or 7 in the skin samples from the Foxn1^**+/+**^ mice (^#^ p<0.05; ^##^p<0.01).

### Skin wound healing in mice is accompanied by an inflammatory/macrophage response

Monocyte chemoattractant protein 1 (MCP-1) is a chemotactic cytokine involved in the induction of lymphocyte and monocyte infiltration to the wound site ^45^. The MCP-1 protein levels in uninjured and injured skin samples showed no differences between the Foxn1^−/−^ and Foxn1^+/+^ mice (Figure 5A, Supplementary Table 12). Next, the content (Figure 5B) and localization (Figure 5C) of CD68-positive macrophages were analysed. Flow cytometric analysis showed that the percentage of CD68-positive cells isolated from the uninjured and injured skin of the Foxn1^−/−^ and Foxn1^+/+^ mice was similar (Figure 5B, Supplementary Table 13). A highly significant increase in the CD68-positive skin cells relative to that of the uninjured (day 0) skin samples was detected at day 5 and 7 post-injury in both the Foxn1^−/−^ (p<0.001) and Foxn1^+/+^ mice (p<0.001; Figure 5B, Supplementary Table 13). The immunohistochemical detection of the CD68-positive cells showed that they were present in the dermis, blood vessels and dermal white adipose tissue (dWAT) after the injury (Figure 5C). The extensive accumulation of CD68-positive cells was observed in the wounded skin at day 5, primarily in the Foxn1^−/−^ mice (Figure 5C).

**Figure 5.**
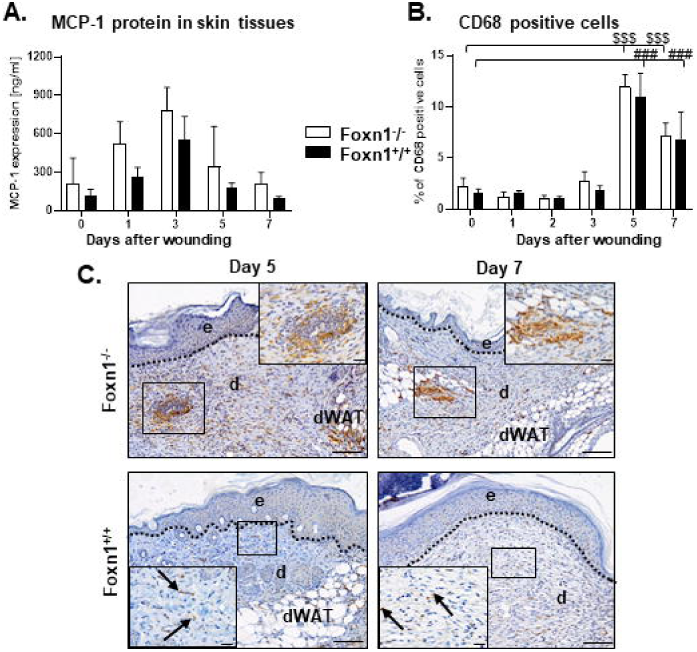
Inflammatory response of the Foxn1^−/−^ and Foxn1^+/+^ mice to excisional skin injury. **(A)** MCP-1 protein levels and (B) flow cytometry analysis of the percentage of CD68^+^ cells isolated from uninjured and injured skin of the Foxn1^**−/−**^ and Foxn1^**+/+**^ mice (n=3–5 per group). (C) Immunohistological localization of CD68 in the skin from the Foxn1^**+/+**^ and Foxn1^**−/−**^ mice collected at days 5 and 7 post-injury. e - epidermis, d-dermis, dWAT-dermal white adipose tissue. Scale bar 100 µm, insets 50 µm. Values are the lsmean ± SE; # indicates significant differences between day 0 vs 5 and 0 vs 7 (^###^p<0.001) in the skin samples of the Foxn1^**+/+**^ mice; $ indicates significant differences between day 0 vs 5 and 0 vs 7 (^$$$^p<0.001) in the skin samples of the Foxn1^**−/−**^ mice.

### The wounded skin of the Foxn1^−/−^ mice displays a distinctive *collagen, Tgfβ* and *Mmp-9/Timp-1* expression profile

Restoration of skin homeostasis and its integrity after injury require biosynthesis of collagen, which is the main component of the ECM ^46^. To evaluate collagen accumulation during skin wound healing, we determined *collagen I* and *III* mRNA expression (Figure 6A, 6B) and the total collagen content through hydroxyproline analysis (Figure 6C), and Masson’s trichrome staining of histological sections of the skin (Figure 6D) was performed.

**Figure 6.**
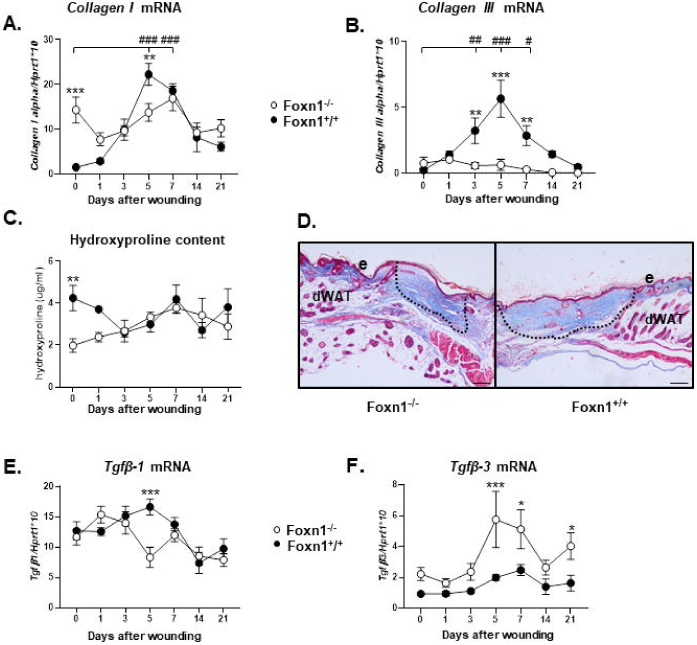
Distinctive expression of collagens and Tgfβs in the skin of the Foxn1^−/−^ mice. qRT-PCR analysis of *collagen I* **(A)**, *collagen III* **(B)**, *Tgfβ-1* **(E)** and *Tgfβ-3* **(F)** mRNA expression during the course of skin wound healing in the Foxn1^−/−^ and Foxn1^+/+^ mice (n=6 per group). **(C)** Hydroxyproline content (n=4 samples per group) in the skin of the Foxn1^−/−^ and Foxn1^**+/+**^ mice. **(D)** Representative histological sections stained with Masson trichrome collected from the skin of the Foxn1^**−/−**^ and Foxn1^**+/+**^ mice at day 21 post-injury. e-epidermis, dWAT-dermal white adipose tissue. Scale bar 200 µm. Values are the lsmean ± SE; asterisks indicate significant differences between the Foxn1^**−/−**^ and Foxn1^**+/+**^ mice during the skin wound healing process (*p < 0.05;** p < 0.01; ***p < 0.001); # indicates significant differences between day 0 and the days of healing in the Foxn1^**+/+**^ mice (^#^p<0.05; ^##^p<0.01; ^###^p<0.001).

The *collagen I* mRNA levels in uninjured (d0) skin were significantly higher in the Foxn1^−/−^ mice than in the Foxn1^+/+^ mice (p<0.001, Figure 6A, Supplementary Tables 14–15). During wound healing, the highest levels of *collagen I* were detected in the skin of the Foxn1^+/+^ mice at day 5 post-injury (Figure 6A, Supplementary Tables 14–15). The analysis of the *collagen III* mRNA expression showed substantial differences between the Foxn1^−/−^ and Foxn1^+/+^ mice (Figure 6B). The uninjured skin of both groups of mice showed similar *collagen III* mRNA levels, but the injured skin of the Foxn1^+/+^ mice displayed a sharp increase in *collagen III* mRNA expression, particularly at days 3 (p<0.01), 5 (p<0.001) and 7 post-wounding (p<0.01), compared to the uninjured control skin (Figure 6B, Supplementary Tables 16–17). In contrast, in the Foxn1^−/−^ mice, the *collagen III* mRNA levels remained stable and unchanged during the entire wound healing process (Figure 6B, Supplementary Tables 16– 17). Total collagen content, measured by hydroxyproline assay, was much lower in the uninjured skin of the Foxn1^−/−^ than in that of the Foxn1^+/+^ mice (p<0.01, Figure 6C, Supplementary Table 18). At days 3–21 post-injury, the hydroxyproline content between the Foxn1^−/−^ and Foxn1^+/+^ mice was comparable (Figure 6C). Masson trichrome-stained skin samples showed the accumulation of collagen fibres in the injured areas in both the Foxn1^−/−^ and Foxn1^+/+^ mice (Figure 6D). TGFβ-1 and TGFβ-3 are cytokines involved in all stages of skin wound healing ^47^. TGFβ-1 is regarded as a profibrotic isoform, whereas TGFβ-3 has antifibrotic functions ^48^. Generally, the uninjured and injured skin of the Foxn1^−/−^ and Foxn1^+/+^ mice showed similar levels of *Tgfβ-1* expression. The only difference was a significant decrease in the expression of *Tgfβ-1* in the Foxn1^−/−^ mice, which was observed at day 5 after wounding (p<0.001; Figure 6E, Supplementary Tables 19–20). qRT-PCR analysis of *Tgfβ-3* gene expression showed significant differences between the Foxn1^−/−^ and Foxn1^+/+^ mice (Figure 6F, Supplementary Tables 21–22). Although the *Tgfβ-3* mRNA expression pattern was similar between the two groups of mice, the magnitude of expression was significantly higher in the Foxn1^−/−^ mice than the Foxn1^+/+^ mice in both uninjured and injured skin. The analysis showed an increase in *Tgfβ-3* expression in the skin of the Foxn1^−/−^ mice at days 5 and 7 (p<0.001, p<0.05, respectively) and day 21 (p<0.05, Figure 6F, Supplementary Table 21) relative to their Foxn1^+/+^ counterparts.

Next, we analysed matrix metalloproteinase 9 (Mmp-9) and its tissue inhibitor (Timp-1), enzymes involved in the inflammatory, proliferative and remodelling phases of wound healing, particularly regulating ECM content ^49^ (Figure 7). Generally, skin injury caused sharp increases in *Mmp*-9 (day 5, p<0.001, Figure 7A, Supplementary Tables 23–24) and *Timp-1* (day 1, p<0.001, Figure 7B, Supplementary Tables 25–26) in the Foxn1^+/+^ mice. The Foxn1^−/−^ mice responded to skin injury with slight, statistically insignificant increases in Mmp-9 (mRNA expression and protein levels; Figure 7A and 7C, respectively) and *Timp-1* mRNA expression (Figure 7B). The Foxn1^+/+^ and Foxn1^−/−^ mice showed major differences in *Mmp-9* mRNA expression (Figure 7A) and Mmp-9 protein levels (Figure 7C) at day 5 post-injury (Supplementary Table 23). Higher levels of *Timp-*1 mRNA expression were detected in the Foxn1^+/+^ mice than the Foxn1^−/−^ mice at days 1–5 post-wounding (Figure 7B, Supplementary Table 25).

**Figure 7.**
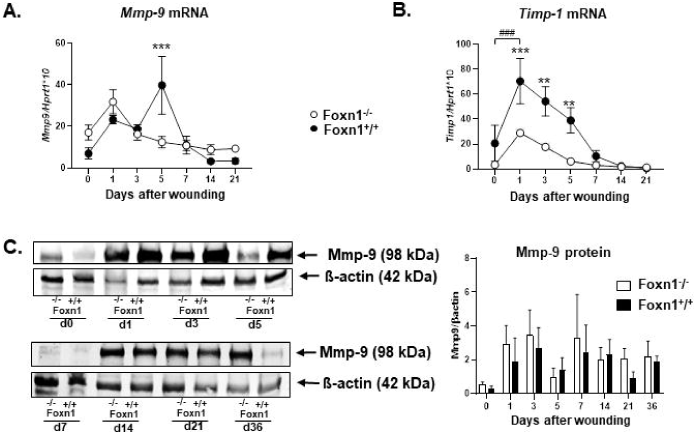
The *Mmp-9/Timp-1* balance in the wounded skin of the Foxn1^−/−^ mice is shifted towards higher *Mmp-9* and lower *Timp-1* levels. *Mmp-9* **(A)** and *Timp-1* **(B)** mRNA expression (n=6 skin samples per group). **(C)** Representative Western blot and densitometric analysis of Mmp-9 in the skin of the Foxn1^**−/−**^ and Foxn1^**+/+**^ mice during wound healing (n=3 skin samples per group). Values are the lsmean ± SE; asterisks indicate significant differences between the Foxn1^**−/−**^ and Foxn1^**+/+**^ mice (**p < 0.01;***p < 0.001); # indicates significant differences between day 0 vs day 1 (^##^p <0.01) in the skin samples from the Foxn1^**+/+**^ mice.

### The effects of TGFβ-1 and TGFβ-3 on dermal fibroblasts (DFs)

DFs phenotype and function are modulated by many growth factors, including TGFβ ^50^. Consistent with the *in vivo* data (see Figures 6E and 6F), which showed the differences in *Tgfβ* mRNA expression in the skin of the Foxn1^−/−^ and Foxn1^+/+^ mice, we next investigated the effect of Tgfβ-1 or Tgfβ-3 on DFs isolated from the Foxn1^−/−^ and Foxn1^+/+^ mice *in vitro* (Figure 8).

**Figure 8.**
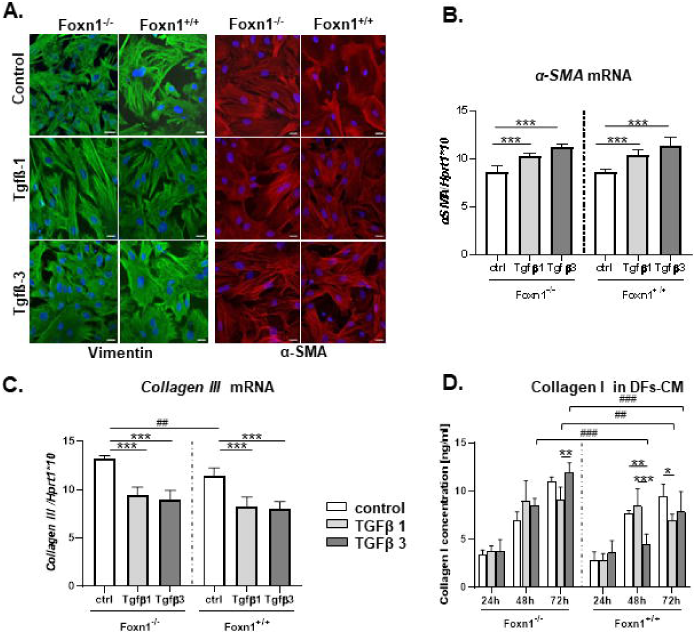
Tgfβ-1 and Tgfβ-3 modulate DF phenotype and collagen production. Representative confocal imaging of vimentin and α-SMA **(A)** in the control and Tgfβ-1- or Tgfβ-3-treated Foxn1^**−/−**^ and Foxn1^**+/+**^ DFs. qRT-PCR analyses of *α-SMA* **(B)** and *collagen III* **(C)** in the control and Tgfβ-1- or Tgfβ-3-treated Foxn1^**−/−**^ and Foxn1^**+/+**^ DFs. Enzyme immunoassay (EIA) analysis of collagen I **(D)** concentration in conditioned media collected from the Foxn1^**−/−**^ and Foxn1^**+/+**^ DFs treated with Tgfβ-1 or Tgfβ-3. Values are the lsmean ± SE; asterisks indicate significant differences between treatments (ctrl vs Tgfβ-1 vs Tgfβ-3) in the Foxn1^**−/−**^ or Foxn1^**+/+**^ DFs (*p < 0.05;** p < 0.01;*** p < 0.001); # indicates significant differences between Foxn1^**−/−**^ vs Foxn1^**+/+**^ by time and condition of culture (^##^p <0.01, ^###^p <0.001).

Immunofluorescence analysis showed the localization of vimentin (marker of fibroblasts) and α-smooth muscle actin (α-SMA; marker of myofibroblasts) in the control and Tgfβ-1- or Tgfβ-3-treated DFs isolated from the Foxn1^−/−^ and Foxn1^+/+^ mice (Figure 8A). Analysis of gene expression by quantitative RT-PCR revealed that Tgfβ-1 or Tgfβ-3 treatment increased the *α-Sma* mRNA expression in both the Foxn1^−/−^ (p<0.001) and Foxn1^+/+^ (p<0.001) cultures compared with the control, untreated cultures (Figure 8B, Supplementary Table 27). The analysis of *collagen* III expression demonstrated that the basal level of its transcript was higher in the DFs from the Foxn1^−/−^ group than the Foxn1^+/+^ group (p<0.01; Figure 8C, Supplementary Tables 28–29). Hence, 24 h treatment with Tgfβ-1 or Tgfβ-3 decreased the expression of *collagen* III in both the Foxn1^−/−^ and Foxn1^+/+^ groups (p<0.001 for both cohorts) relative to that of the untreated DFs and led to comparable levels between the Foxn1^−/−^ and Foxn1^+/+^ DFs (Figure 8C, Supplementary Tables 28–29).

One of the main functions of DFs is the production of ECM components, including collagen I, a major skin protein ^51^. To examine the potential differences between DFs from the Foxn1^−/−^ and Foxn1^+/+^ mice, we cultured both types of DFs in monolayers and treated them with Tgfβ-1 or Tgfβ-3 for 24, 48 and 72 h. Conditioned media were collected at each time point. Enzyme immunoassays (EIAs) revealed that the concentration of collagen I in the conditioned media increased with the time of culture (24 h<48 h<72 h), regardless of whether Foxn1^−/−^ or Foxn1^+/+^ DFs were used (Figure 8D; Supplementary Table 30–32).

Generally, at 48 and 72 h of treatment, the collagen I concentration was higher in the conditioned media collected from the Foxn1^−/−^ DFs than from the Foxn1^+/+^ DFs, especially for the Tgfβ-3-treated DFs (p<0.001 for both 48 h and 72 h; Figure 8D, Supplementary Table 30– 32). Interestingly, Tgfβ-3 (at 48 h) and Tgfβ-1 (at 72 h) had inhibitory effects on collagen I secretion in the DFs isolated from the skin of the Foxn1^+/+^ mice compared to the untreated controls (p<0.01 for Tgfβ-3 and p<0.05 for Tgfβ-1; Figure 8D, Supplementary Tables 30–32).

### Effects of hypoxia on the expression of Foxn1 and Hif-1α in regenerative (Foxn1^−/−^) and reparative (Foxn1^+/+^) keratinocytes

Our recent *in vitro* study showed that the expression of Foxn1 is induced by hypoxia ^23^. Moreover, the present *in vivo* data (see Figure 1 and 2) revealed low levels of hypoxia-regulated factors (Hif-1α and Trx-1) in the skin of the Foxn1^−/−^ mice. Therefore, we next investigated how oxygen availability (21% O_2_ vs 1% O_2_) and time of culture (12 h vs 24 h) affected the expression of *Foxn1, Hif-1a* and *Fih-1* in keratinocytes isolated from the Foxn1^+/+^ (Figures 9A, 9C, 9E) or Foxn1^−/−^ mice (Figures 9B, 9D, 9F) that were cocultured with DFs. The Foxn1^−/−^ keratinocytes were transduced with Ad-Foxn1 or Ad-GFP (control) vectors prior to coculture with DFs and exposure to different levels of oxygen.

**Figure 9.**
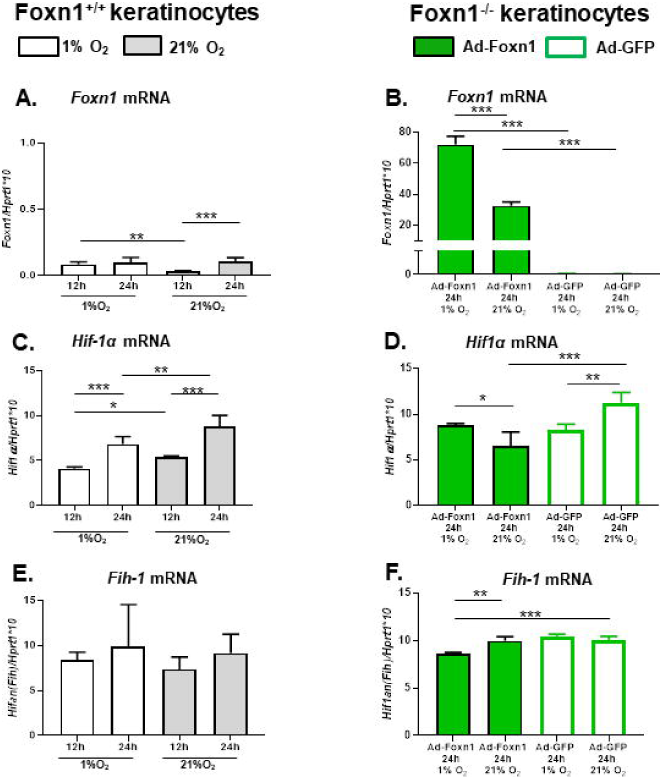
Foxn1 modulates *Hif-1α* and *Fih-1* expression in keratinocytes cultured in normoxic vs hypoxic microenvironments. Keratinocytes isolated from the Foxn1^+/+^ **(A, C, E)** or Foxn1^**−/−**^ **(B, D, F)** mice were cocultured with DFs in normoxic (21% O_2_) or hypoxic (1% O_2_) conditions. mRNA expression levels of *Foxn1* **(A, B)**, *Hif-1α* **(C, D)** and *Fih-1* **(E, F)** were analysed. Keratinocytes isolated from the Foxn1^−/−^ mice (**B, D, F**) prior to culture in 21% vs 1% O_2_ were transduced with the Ad-Foxn1 or Ad-GFP (control) vectors. Values are the lsmean ± SE; asterisks indicate significant differences between normoxic and hypoxic conditions or time of culture (*p < 0.05;** p < 0.01;*** p < 0.001).

In the Foxn1^+/+^ keratinocytes, hypoxic (1% O_2_ oxygen availability) but not normoxic (21% O_2_) conditions increased endogenous Foxn1 expression during the first 12 h of culture (p<0.01) to levels that were maintained for the next 12 h of culture (Figure 9A, Supplementary Table 34). Interestingly, under normoxia, the low *Foxn1* mRNA level at 12 h of culture significantly increased at 24 h (p<0.001, Figure 9A, Supplementary Table 33) to reach a comparable level to that of the cultures maintained in normoxic or hypoxic conditions. In the Foxn1^−/−^ keratinocytes that were transduced with Ad-Foxn1, hypoxia (1% O_2_) strongly stimulated the exogenous *Foxn1* mRNA levels compared to the levels expressed under normoxic conditions (p<0.001) (Figure 9B, Supplementary Tables 35–36). Next, using the same experimental settings, i.e., Foxn1^+/+^ or Foxn1^−/−^ keratinocytes cultured with different oxygen availability, we analysed *Hif-1α* mRNA expression (Figures 9C and 9D). The Foxn1^+/+^ keratinocytes showed an increase in *Hif-1α* mRNA expression during culture time (12 h<24 h) regardless of the culture conditions: normoxia or hypoxia (Figure 9C).

Interestingly, the increase in the *Hif-1α* mRNA levels over time was higher in normoxic than hypoxic (1% O_2_) environments regardless of culture time (p<0.001 for both 12 h or 24 h; Figure 9C, Supplementary Tables 37–38). No differences in *Fih-1* mRNA expression regardless of culture time or oxygen availability were detected in the Foxn1^+/+^ keratinocytes (Figure 9E).

The keratinocytes deficient in endogenous Foxn1 (Foxn1^−/−^) that were transduced with Ad-Foxn1 (exogenous Foxn1) showed different responses than the Foxn1^+/+^ keratinocytes to culture conditions with respect to the expression of the hypoxia-regulated factors *Hif-1α* (Figure 9D) and *Fih-1* (Figure 9F). First, control keratinocytes that were transduced with Ad-GFP and lacked endogenous or exogenous Foxn1 showed an increase in *Hif-1α* mRNA under normoxic conditions compared with hypoxic conditions (p<0.01), similar to the Foxn1^+/+^ keratinocytes (compare Figures 9C and 9D, Supplementary Tables 39–42). In contrast, the Ad-Foxn1-transduced keratinocytes showed a significant reduction in *Hif-1α* mRNA expression in normoxic relative to hypoxic cultures (p<0.05; Figure 9D, Supplementary Tables 39–42). Moreover, the mRNA levels of factors inhibiting Hif-1 (*Fih-1, Hif-1an*) were inhibited by hypoxia (p<0.01) exclusively in the Ad-Foxn1 transduced keratinocytes and not in the control-Ad-GFP-transduced cells (Figure 9F, Supplementary Tables 43–44).

## DISCUSSION

The aim of the present study was to investigate the potential correlation between Foxn1 and hypoxia-regulated elements (Hif-1α, Fih-1 and Trx-1) and its impact on the skin wound healing process. Using *in vivo* and *in vitro* approaches, we identified the transcription factor Foxn1 as a potential regulator of Hif-1α stability during cutaneous wound healing. The data indicated that the interplay between Foxn1 and hypoxia-regulated factors may direct the outcome of cutaneous repair: the reparative (scar-forming) outcomes observed in the Foxn1^+/+^ mice vs the regenerative (scarless) outcomes in the Foxn1^−/−^ mice.

The present *in vivo* study performed on an excisional model of skin injury confirmed our previous data on an incisional model and showed that the Foxn1^−/−^ mice displayed a notable ability for scarless (regenerative) healing ^10, 13, 16^. Although the incisional and excisional wounds showed different healing profiles ^52^, we observed that both types of skin trauma (incisional vs excisional) shared similar scarless healing patterns in the Foxn1^−/−^ mice. Indeed, excisionally wounded skin of the Foxn1^−/−^ mice revealed several features attributed to the regenerative healing of incisional wounds, including fast re-epithelialization, increased *Tgfβ-3* (anti-scarring) cytokine expression that correlated with a decrease in *Tgfβ-1* (pro-scarring) expression, and a shift in the *Mmp-9*/*Timp-1* balance towards higher *Mmp-9* and lower *Timp-1* levels ^10, 13, 16^. However, some differences between incisional and excisional skin injury were observed due to the amount of collagen deposition in the wounded skin of the Foxn1^−/−^ mice. The injured skin of incisional wounds of the Foxn1^−/−^ mice at day 21 displayed a narrow almost undetectable line of collagen bundles, barely indicating the place of injury ^13^. The skin of the excisional wounds of the Foxn1^−/−^ mice at day 21 post-injury showed good demarcation from the surrounding tissue deposition of new collagen, although the area of collagen accumulation was smaller than that in the Foxn1^+/+^ mice. These particular data support the concept that the type of injury and amount of trauma reflect the healing profile ^52^.

Notably, the present data indicate that hypoxia-regulated factors are controlled through Foxn1. The injured skin of the Foxn1^−/−^ mice showed stable and unchanged expression of *Hif-1α* and *Fih-1*, the key elements of tissues that sense oxygen deprivation. In contrast, the skin of the Foxn1^+/+^ mice displayed an increase in *Hif-1α* and a decrease in *Fih-1* in response to excisional wounds. Since the skin of the Foxn1^−/−^ mice healed in a scarless manner, the data suggest that impairment in Hif-1α due to the lack of Foxn1 redirects the healing pathway from repair (scar-present) towards regeneration (scarless). The *in vitro* approach revealed that Foxn1 overexpression in keratinocytes isolated from the skin of Foxn1^−/−^ mice led to reduced *Hif-1α* mRNA expression in normoxic but not hypoxic cultures and inhibited *Fih-1* expression exclusively under hypoxic conditions.

Extensive data have demonstrated that Hif-1α, a master regulator of oxygen homeostasis, participates in tissue repair ^3, 30, 53^. Moreover, an imbalance in Hif-1α expression is associated with adverse skin wound healing outcomes. Previous studies showed that overexpression of Hif-1α leads to fibrotic disease, whereas Hif-1α deficiency results in chronic hypoxia, which contributes to the formation of nonhealing wounds ^3, 30, 32, 39, 54^. Interestingly, scar-free skin wound healing observed in mammalian foetuses occurs in a physiologically hypoxic environment ^41^. Moreover, differences in Hif-1α expression between the regenerative healing of foetal skin and the reparative healing of adult skin have been detected ^55^. Scheid et al., showed no Hif-1α expression in the wounded skin of sheep foetuses (scar-free period of healing) and increased levels of Hif-1α in the skin of adult sheep during scar-forming healing ^55^. The results of our investigations are consistent with the study by Scheid et al.,^41^. The Foxn1^−/−^ mice, which heal skin injuries in a scarless manner, displayed low and unchanged levels of Hif-1α in the healing process. In addition, the injured skin tissues of the Foxn1^−/−^ mice showed low and stable expression of Fih-1 during the healing process.

The present *in vivo* and *in vitro* data also identified factors regulated by hypoxia that are linked with Foxn1. The Trx-1 levels on days 3 and 7 were higher in the wounded skin of the Foxn1^+/+^ mice than that of the Foxn1^−/−^ mice. Furthermore, double immunofluorescence staining showed colocalization of Trx-1 with Foxn1 in the epidermis, particularly in the suprabasal layer. Shaikh et al., revealed that Hif-1α activity depends on Trx-1 redox status ^56^. Trx-1 scavenges reactive oxygen species during oxidative stress and rescues injured cells from hypoxia to maintain homeostasis ^57^. As reported by Naranjo-Suarez et al., overexpression of Trx-1 in HeLa, HT-29, MCF-7 and EMT6 cell lines did not stabilize Hif-1α but led to increased activity ^58^. Other studies have shown that Trx-1 acts as a transcription factor that amplifies Hif-1α synthesis ^34, 59^.

Hypoxic conditions induce not only *Hif-1α* but also TGFβ, as observed in cancer cells ^60^. In the skin, TGFβ-1 and TGFβ-3 modulate cutaneous wound healing. Whereas TGFβ-1 is regarded as a profibrotic isoform, TGFβ-3 has an antifibrotic function ^48, 61^. A study by Whitby and Ferguson showed that scarless healing in mammalian embryos is due to elevated levels of TGFβ-3 ^62^. Furthermore, Shah et al., revealed that TGFβ-3 administration caused a reduction in scar formation in adult rats ^63^. Consistently, our current *in vivo* study revealed that skin from the Foxn1^−/−^ mice exhibited significantly higher mRNA levels of *Tgfβ-3* than that of the Foxn1^+/+^ mice. Interestingly, high levels of *Tgfβ-3* expression in the skin of the Foxn1^−/−^ mice at day 5 post-wounding were accompanied by low levels of *Tgfβ-1*. This result suggests that *Tgfβ-3* contributes to the regenerative capacity of the Foxn1^−/−^ mice; however, the possible mechanisms of interaction among Foxn1, hypoxia and Tgfβ require further investigation. Considering that the primary mechanism of healing is associated with the deposition of ECM proteins, we measured the concentration of collagen I produced by DFs treated with Tgfβ-1 or Tgfβ-3. The results showed a higher concentration of collagen type I in conditioned media collected from the Foxn1^−/−^ DFs than from the Foxn1^+/+^ DFs, particularly in the Tgfβ-3-treated DFs. Similar results were obtained by Bukowska et al., showing that the content of collagen type I in culture media collected from Foxn1^−/−^ DFs increased following TGFβ-3 but not TGFβ-1 administration ^64^. Collectively, the present data showed that DFs originating from the Foxn1^−/−^ mice are more sensitive to stimulation with Tgfβ-3 than those isolated from the Foxn1^+/+^ mice.

The mechanisms that underlie the differences between reparative and regenerative healing are still under debate. Our *in vivo* data showed that Foxn1 and hypoxia-regulated factors (Hif-1α, Fih-1, Trx-1) expressed in cutaneous epithelia, particularly in the epidermal suprabasal layer, regulate the skin wound healing process. To elucidate the possible mechanism of interaction between Foxn1 and hypoxia-related factors, we compared how oxygen availability and time of culture affect the expression of *Foxn1, Hif-1α* and *Fih-1* in keratinocytes isolated from the Foxn1^+/+^ or Foxn1^−/−^ mice. First, we confirmed our previous data showing that hypoxic conditions stimulated Foxn1 expression ^23^. Next, we showed that Foxn1 transduction in Foxn1^−/−^ keratinocytes downregulated *Hif-1α* and upregulated *Fih-1* in normoxic but not hypoxic conditions. The data suggest that under normoxic conditions, Foxn1 is a limiting factor for Hif-1α. Similar findings were demonstrated by Wang et al., using an RNA silencing of nuclear respiratory factor-1 (NRF-1) ^65^. Depletion of NRF-1 in HEK293T cells increased Hif-1α protein accumulation under normoxia, and this effect was further amplified under hypoxia^65^.

A large body of data has shown that transcription factors such as heat shock factors HSF2 and HSP4 and early growth response 1 (Egr-1) upregulate Hif-1α under hypoxia ^66 67^. Moreover, regulation of Hif-1α transcription was observed in cancerous tissues; for example, in hepatocellular carcinoma (HCC), Bclaf-1 enhances transcription of Hif-1α ^68^. In contrast, signal transducer and activator of transcription 3 (Stat3) inhibits the oncogenic potential of HIF-1α induced by hypoxia ^69^. Other studies have demonstrated that some non-hypoxic stimuli can enhance Hif-1α, e.g., thrombin and angiotensin II ^70, 71^. Our *in vivo* and *in vitro* data showed that Foxn1 modulates the hypoxic regulatory pathway involving Hif-1α and Fih-1. Overexpression of *Foxn1* (Ad-Foxn1) altered the balance between *Hif-1α* and *Fih*-1 under normoxic conditions. Indeed, the abundance of Foxn1 led to downregulation of *Hif-1α* mRNA and upregulation of the *Fih-1* transcript. Shifting the balance between *Hif-1α* and *Fih-1* by Foxn1 introduces possible new mechanisms of inactivation of the cell survival pathway in the skin that are unnecessary under normoxia. The mechanism by which Foxn1 reduces Hif-1α gene transcription is possibly achieved through PKC signalling. Our previous study confirmed that Foxn1 is a negative regulator of diacylglycerol-sensitive protein kinase C (PKC) activity in keratinocytes ^21, 23^. The activation of PKC was shown to play a major role in the increase in Hif-1α gene transcription ^72^. Xia et al., analysed the effect of vitamin K2 (VK2) on HCC cells and revealed that VK2 can suppress HIF-1α activation by inhibiting PKC in HCC cells ^73^. Keratinocytes synthesize up to five PKC isoforms, among which PKCα, δ, ε, and η exhibited higher levels in keratinocytes from the Foxn1^−/−^ than Foxn1^+/+^ mice ^21^. Li et al., showed that Foxn1 inhibits the activation of PKCδ, the isoform whose inhibition in HCC cells suppresses HIF-1α activation ^73^. This finding led to the suggestion that Foxn1 downregulates PKC signalling in keratinocytes, which in turn regulates Hif-1α transcription. Further experiments are required to verify the aforementioned concept.

Our previous study revealed that Foxn1 is one of the key regulators in the skin wound healing process ^28, 29, 42^. The expression of Foxn1 in the skin is limited to the epidermis and hair follicles. The activity of Foxn1 during wound healing is associated with epidermal reconstruction (re-epithelialization) and facilitates scar formation due to its involvement in EMT ^28, 29^. In addition, recent data showed that Foxn1 regulates the adipogenic potential of the skin (dermal white adipose tissues; dWAT), thereby contributing to the skin wound healing process ^42^. The present data suggest that Foxn1 may modulate skin homeostasis (keratinocyte proliferation/differentiation) and the skin response to wounding through the regulation of the skin response to different (normoxia vs hypoxia) levels of oxygenation. However, the precise mechanisms by which Foxn1 regulates the skin wound healing process are still unclear and await clarification.

In conclusion, our data suggest that the transcription factor Foxn1 (Foxn1^+/+^ mice), as a potential regulator of Hif-1α, governs the reparative (scar-present) skin wound healing process. In contrast, Foxn1-deficient (Foxn1^−/−^ mice) mice exhibited Hif-1α deregulation that resulted in regenerative (scarless) skin wound healing. The data further underline Foxn1 as a key element of the maintenance of skin homeostasis. Moreover, we propose that the changes in the expression of wound healing-associated factors (hypoxia-related genes, Foxn1, Mmps/Timps, collagen) position scar-free healing (regeneration) between scar formation and nonhealing wounds (Figure 10). The subtle differences in the timing and quantity of expression of “post-wounding effector genes” may alter the proportions among reparative, regenerative and non-healing pathways. Therefore, the present study underscores the role of Foxn1 in oxygen homeostasis during the skin healing process. The link between Foxn1 and hypoxia-related factors may have an impact on wound healing resolutions such as regeneration or scar formation or, alternatively, its dysfunction might result in an inability to heal.

**Figure 10.**
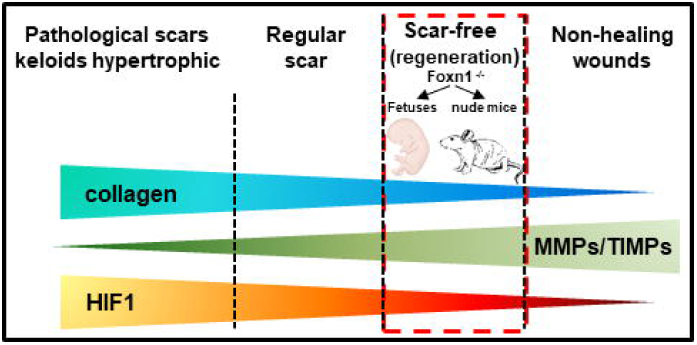
Changes in wound healing-associated factors (hypoxia-related genes, Foxn1, Mmps/Timps, collagen) position scar-free healing (regeneration) between scar-forming healing and nonhealing wounds.

## ACKNOWLEDGEMENTS

The research in the Gawronska-Kozak laboratory is supported by the National Science Centre, Poland; Grant OPUS 14 No.2017/27/B/NZ5/02610. We are grateful to The Support Staff of The Center of Experimental Medicine (CEM), Medical University of Bialystok for assistance in animal study.

## AUTHOR CONTRIBUTIONS

Designed research: B. Gawronska-Kozak; Performed research: S. Machcińska, M. Kopcewicz, J. Bukowska, K. Walendzik, B. Gawronska-Kozak; Analyzed data: S. Machcińska, B. Gawronska-Kozak, M. Kopcewicz, J. Bukowska, K. Walendzik; Wrote the paper: S. Machcińska, B. Gawronska-Kozak.

