## Supplementary material for "Impairment of the Hif-1α regulatory pathway in Foxn1-deficient (Foxn1^−/−^) mice affects the skin wound healing process": Suppl. files

### Supplementary Figure 1

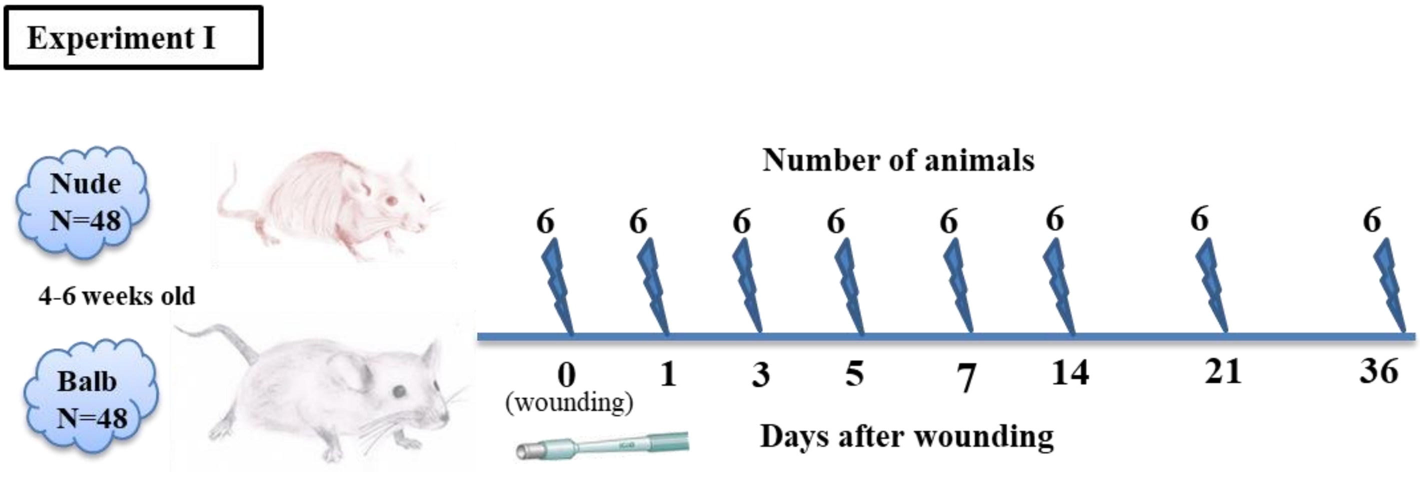


### Supplementary Figure 2

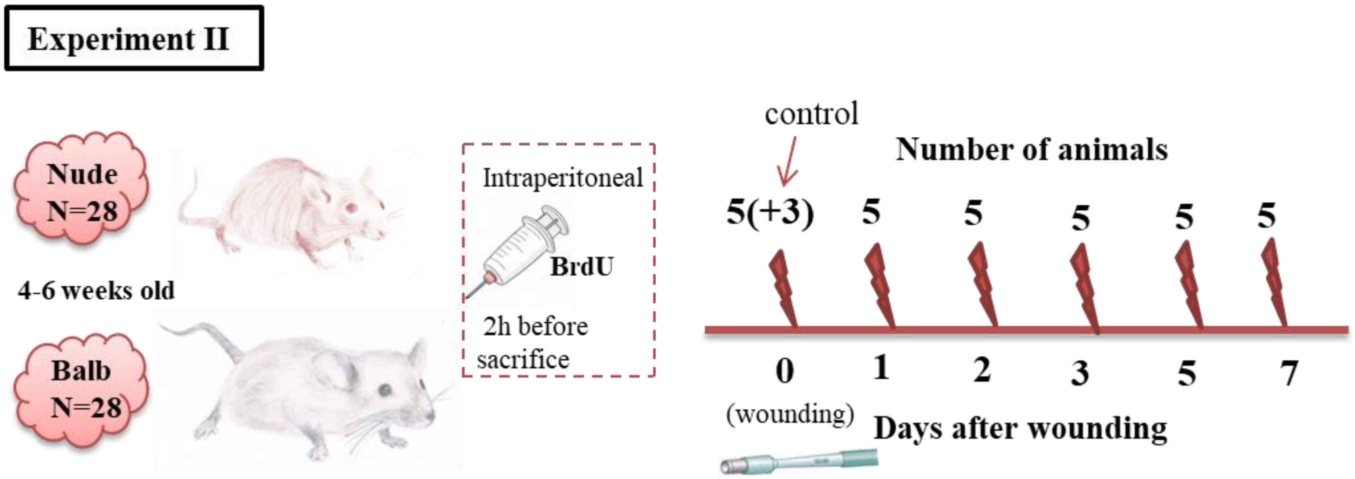

**Statistically significant differences for *Hif-1α* and *Fih-1* expression in Foxn1$^{-/-}$ (CBy.Cg-Foxn1<nu>/cmdb ) and Foxn1$^{+/+}$ (Balb/c/cmdb) mice during wound healing process**

### *Hif-1 α*

**Supplementary Table 1. Comparisons by group for *Hif-1 α***

| Day | group | lsmean | lower.CL | upper.CL | SE | df | p.value |
|---|---|---|---|---|---|---|---|
| Day 0 | Foxn1$^{+/+}$ | 23.65 | 19.67 | 27.63 | 2.00 | 69.00 | <0.001 |
| Day 0 | Foxn1$^{-/-}$ | 13.66 | 9.68 | 17.64 | 2.00 | 69.00 | |
| Day 1 | Foxn1$^{+/+}$ | 30.97 | 26.99 | 34.96 | 2.00 | 69.00 | <0.001 |
| Day 1 | Foxn1$^{-/-}$ | 20.24 | 16.26 | 24.22 | 2.00 | 69.00 | |
| Day 3 | Foxn1$^{+/+}$ | 28.23 | 23.87 | 32.60 | 2.19 | 69.00 | 0.002 |
| Day 3 | Foxn1$^{-/-}$ | 18.93 | 14.95 | 22.91 | 2.00 | 69.00 | |
| Day 5 | Foxn1$^{+/+}$ | 25.49 | 21.50 | 29.47 | 2.00 | 69.00 | <0.001 |
| Day 5 | Foxn1$^{-/-}$ | 15.66 | 11.68 | 19.65 | 2.00 | 69.00 | |
| Day 21 | Foxn1$^{+/+}$ | 12.28 | 8.30 | 16.26 | 2.00 | 69.00 | 0.049 |
| Day 21 | Foxn1$^{-/-}$ | 17.94 | 13.96 | 21.93 | 2.00 | 69.00 | |

**Supplementary Table 2. Comparisons by days for *Hif-1 α***

| Day | group | lsmean | lower.CL | upper.CL | SE | df | Day 1 | Day 3 | Day 5 | Day 7 | Day 14 | Day 21 |
|---|---|---|---|---|---|---|---|---|---|---|---|---|
| Day 0 | Foxn1$^{+/+}$ | 23.65 | 19.67 | 27.63 | 2.00 | 69.00 | 0.143 | 0.714 | 0.995 | 0.320 | 0.009 | 0.003 |
| Day 1 | Foxn1$^{+/+}$ | 30.97 | 26.99 | 34.96 | 2.00 | 69.00 | | 0.967 | 0.459 | <0.001 | <0.001 | <0.001 |
| Day 3 | Foxn1$^{+/+}$ | 28.23 | 23.87 | 32.60 | 2.19 | 69.00 | | | 0.967 | 0.009 | <0.001 | <0.001 |
| Day 5 | Foxn1$^{+/+}$ | 25.49 | 21.50 | 29.47 | 2.00 | 69.00 | | | | 0.084 | 0.001 | <0.001 |
| Day 7 | Foxn1$^{+/+}$ | 17.50 | 13.51 | 21.48 | 2.00 | 69.00 | | | | | 0.765 | 0.521 |
| Day 14 | Foxn1$^{+/+}$ | 13.37 | 9.38 | 17.35 | 2.00 | 69.00 | | | | | | 1.000 |
| Day 21 | Foxn1$^{+/+}$ | 12.28 | 8.30 | 16.26 | 2.00 | 69.00 | | | | | | |
| Day 0 | Foxn1$^{-/-}$ | 13.66 | 9.68 | 17.64 | 2.00 | 69.00 | 0.245 | 0.509 | 0.992 | 0.999 | 0.999 | 0.733 |
| Day 1 | Foxn1$^{-/-}$ | 20.24 | 16.26 | 24.22 | 2.00 | 69.00 | | 0.999 | 0.669 | 0.542 | 0.527 | 0.983 |
| Day 3 | Foxn1$^{-/-}$ | 18.93 | 14.95 | 22.91 | 2.00 | 69.00 | | | 0.907 | 0.824 | 0.813 | 1.000 |
| Day 5 | Foxn1$^{-/-}$ | 15.66 | 11.68 | 19.65 | 2.00 | 69.00 | | | | 1.000 | 1.000 | 0.983 |
| Day 7 | Foxn1$^{-/-}$ | 15.11 | 11.13 | 19.09 | 2.00 | 69.00 | | | | | 1.000 | 0.952 |
| Day 14 | Foxn1$^{-/-}$ | 15.05 | 11.07 | 19.03 | 2.00 | 69.00 | | | | | | 0.946 |
| Day 21 | Foxn1$^{-/-}$ | 17.94 | 13.96 | 21.93 | 2.00 | 69.00 | | | | | | |

### *Fih-1 (Hif1an)*

**Supplementary Table 3. Comparisons by group for *Fih -1(Hif1-an)***

| Day | group | lsmean | lower.CL | upper.CL | SE | df | p.value |
|---|---|---|---|---|---|---|---|
| Day 1 | Foxn1$^{+/+}$ | 11.91 | 8.19 | 15.62 | 1.86 | 69.00 | 0.019 |
| Day 1 | Foxn1$^{-/-}$ | 18.24 | 14.52 | 21.95 | 1.86 | 69.00 | |
| Day 5 | Foxn1$^{+/+}$ | 11.66 | 7.95 | 15.38 | 1.86 | 69.00 | 0.005 |
| Day 5 | Foxn1$^{-/-}$ | 19.36 | 15.65 | 23.08 | 1.86 | 69.00 | |

**Statistically significant differences for: Wound size and % of reepithelization in Foxn1 $^{-/-}$ ( CBy.Cg-Foxn1<nu>/cmdb ) and Foxn1$^{+/+}$ ( Balb/c/cmdb) mice during wound healing process**

### Wound size

**Supplementary Table 4. Comparisons by group for Wound size**

| Day | group | lsmean | lower.CL | upper.CL | SE | df | p.value |
|---|---|---|---|---|---|---|---|
| Day 3 | Foxn1$^{-/-}$ | 7.40 | 3.59 | 11.20 | 1.89 | 50.00 | 0.007 |
| Day 3 | Foxn1$^{+/+}$ | 14.94 | 11.13 | 18.74 | 1.89 | 50.00 | |

**Supplementary Table 5. Comparisons by days for Wound size**

| Day | group | lsmean | lower.CL | upper.CL | SE | df | Day 1 | Day 3 | Day 5 | Day 7 |
|---|---|---|---|---|---|---|---|---|---|---|

| Day | group | lsmean | lower.CL | upper.CL | SE | df | Day 1 | Day 3 | Day 5 | Day 7 |
|---|---|---|---|---|---|---|---|---|---|---|
| Day 0 | Foxn1$^{-/-}$ | 12.57 | 8.76 | 16.37 | 1.89 | 50.00 | 0.441 | 0.314 | 0.026 | 0.003 |
| Day 1 | Foxn1$^{-/-}$ | 8.01 | 4.21 | 11.81 | 1.89 | 50.00 | | 0.999 | 0.642 | 0.195 |
| Day 3 | Foxn1$^{-/-}$ | 7.40 | 3.59 | 11.20 | 1.89 | 50.00 | | | 0.778 | 0.293 |
| Day 5 | Foxn1$^{-/-}$ | 4.31 | 0.51 | 8.11 | 1.89 | 50.00 | | | | 0.922 |
| Day 7 | Foxn1$^{-/-}$ | 2.11 | -1.69 | 5.91 | 1.89 | 50.00 | | | | |
| Day 0 | Foxn1$^{+/+}$ | 12.57 | 8.76 | 16.37 | 1.89 | 50.00 | 1.000 | 0.901 | 0.071 | 0.004 |
| Day 1 | Foxn1$^{+/+}$ | 13.12 | 9.32 | 16.92 | 1.89 | 50.00 | | 0.960 | 0.043 | 0.002 |
| Day 3 | Foxn1$^{+/+}$ | 14.94 | 11.13 | 18.74 | 1.89 | 50.00 | | | 0.007 | <0.001 |
| Day 5 | Foxn1$^{+/+}$ | 5.38 | 1.58 | 9.18 | 1.89 | 50.00 | | | | 0.808 |
| Day 7 | Foxn1$^{+/+}$ | 2.45 | -1.36 | 6.25 | 1.89 | 50.00 | | | | |

### % of re-epithelization

**Supplementary Table 6. Comparisons by group for % of re-epithelization**

| Day | group | lsmean | lower.CL | upper.CL | SE | df | p.value |
|---|---|---|---|---|---|---|---|
| Day 2 | Foxn1$^{+/+}$ | 20.89 | 0.42 | 41.36 | 9.66 | 16.00 | 0.002 |
| Day 2 | Foxn1$^{-/-}$ | 70.32 | 49.86 | 90.79 | 9.66 | 16.00 | |

**Supplementary Table 7. Comparisons by days for % of re-epithelization**

| Day | group | lsmean | lower.CL | upper.CL | SE | df | Day 2 | Day 3 | Day 5 |
|---|---|---|---|---|---|---|---|---|---|
| Day 1 | Foxn1$^{+/+}$ | 6.35 | -14.12 | 26.81 | 9.66 | 16.00 | 0.715 | <0.001 | <0.001 |
| Day 2 | Foxn1$^{+/+}$ | 20.89 | 0.42 | 41.36 | 9.66 | 16.00 | | 0.002 | <0.001 |
| Day 3 | Foxn1$^{+/+}$ | 80.91 | 60.44 | 101.38 | 9.66 | 16.00 | | | 0.518 |
| Day 5 | Foxn1$^{+/+}$ | 100.00 | 79.53 | 120.47 | 9.66 | 16.00 | | | |
| Day 1 | Foxn1$^{-/-}$ | 24.10 | 3.63 | 44.56 | 9.66 | 16.00 | 0.018 | <0.001 | <0.001 |
| Day 2 | Foxn1$^{-/-}$ | 70.32 | 49.86 | 90.79 | 9.66 | 16.00 | | 0.333 | 0.173 |
| Day 3 | Foxn1$^{-/-}$ | 94.19 | 73.72 | 114.66 | 9.66 | 16.00 | | | 0.973 |
| Day 5 | Foxn1$^{-/-}$ | 100.00 | 79.53 | 120.47 | 9.66 | 16.00 | | | |

### Statistically significant differences for % of BrdU, Cytokeratin 6 (CK6) or CD68 positive cells and MCP-1 protein content isolated from the skin of Foxn1$^{-/-}$ (CBy.Cg-Foxn1<nu>/cmdb) and Foxn1$^{+/+}$(Balb/c/cmdb) mice during wound healing process

### BrdU

**Supplementary Table 8. Comparisons by group for BrdU**

| Day | group | lsmean | lower.CL | upper.CL | SE | df | p.value |
|---|---|---|---|---|---|---|---|
| Day 2 | Foxn1$^{+/+}$ | 2.40 | 1.23 | 3.57 | 0.58 | 44.00 | 0.003 |
| Day 2 | Foxn1$^{-/-}$ | 4.94 | 3.77 | 6.11 | 0.58 | 44.00 | |

**Supplementary Table 9. Comparisons by days for BrdU**

| Day | group | lsmean | lower.CL | upper.CL | SE | df | Day 1 | Day 2 | Day 3 | Day 5 | Day 7 |
|---|---|---|---|---|---|---|---|---|---|---|---|
| Day 0 | Foxn1$^{+/+}$ | 2.80 | 1.29 | 4.31 | 0.75 | 44.00 | 0.871 | 0.998 | 0.162 | 0.879 | 0.998 |
| Day 1 | Foxn1$^{+/+}$ | 1.74 | 0.57 | 2.91 | 0.58 | 44.00 | | 0.965 | 0.643 | 1.000 | 0.473 |
| Day 2 | Foxn1$^{+/+}$ | 2.40 | 1.23 | 3.57 | 0.58 | 44.00 | | | 0.200 | 0.969 | 0.915 |
| Day 3 | Foxn1$^{+/+}$ | 0.48 | -0.69 | 1.65 | 0.58 | 44.00 | | | | 0.628 | 0.020 |
| Day 5 | Foxn1$^{+/+}$ | 1.76 | 0.59 | 2.93 | 0.58 | 44.00 | | | | | 0.488 |
| Day 7 | Foxn1$^{+/+}$ | 3.22 | 2.05 | 4.39 | 0.58 | 44.00 | | | | | |
| Day 0 | Foxn1$^{-/-}$ | 2.83 | 1.32 | 4.34 | 0.75 | 44.00 | 0.999 | 0.247 | 0.963 | 0.995 | 0.984 |
| Day 1 | Foxn1$^{-/-}$ | 3.14 | 1.97 | 4.31 | 0.58 | 44.00 | | 0.261 | 0.774 | 0.923 | 0.859 |
| Day 2 | Foxn1$^{-/-}$ | 4.94 | 3.77 | 6.11 | 0.58 | 44.00 | | | 0.013 | 0.031 | 0.020 |
| Day 3 | Foxn1$^{-/-}$ | 2.06 | 0.89 | 3.23 | 0.58 | 44.00 | | | | 0.999 | 1.000 |
| Day 5 | Foxn1$^{-/-}$ | 2.34 | 1.17 | 3.51 | 0.58 | 44.00 | | | | | 1.000 |
| Day 7 | Foxn1$^{-/-}$ | 2.20 | 1.03 | 3.37 | 0.58 | 44.00 | | | | | |

### Cytokeratin 6 (CK6)

**Supplementary Table 10. Comparisons by group for CK6**

| Day | group | lsmean | lower.CL | upper.CL | SE | df | p.value |
|---|---|---|---|---|---|---|---|
| Day 2 | Foxn1$^{+/+}$ | 0.52 | 0.26 | 0.78 | 0.13 | 41.00 | 0.033 |
| Day 2 | Foxn1$^{-/-}$ | 0.10 | -0.19 | 0.39 | 0.14 | 41.00 | |
| Day 3 | Foxn1$^{+/+}$ | 0.84 | 0.58 | 1.10 | 0.13 | 41.00 | 0.002 |
| Day 3 | Foxn1$^{-/-}$ | 0.24 | -0.02 | 0.50 | 0.13 | 41.00 | |

**Supplementary Table 11. Comparisons by days for CK6**

| Day | group | lsmean | lower.CL | upper.CL | SE | df | Day 1 | Day 2 | Day 3 | Day 5 | Day 7 |
|---|---|---|---|---|---|---|---|---|---|---|---|
| Day 0 | Foxn1$^{+/+}$ | 0.33 | 0.00 | 0.66 | 0.16 | 41.00 | 0.977 | 0.944 | 0.166 | 0.999 | 0.824 |
| Day 1 | Foxn1$^{+/+}$ | 0.18 | -0.11 | 0.46 | 0.14 | 41.00 | | 0.470 | 0.014 | 0.999 | 0.996 |
| Day 2 | Foxn1$^{+/+}$ | 0.52 | 0.26 | 0.78 | 0.13 | 41.00 | | | 0.489 | 0.717 | 0.164 |
| Day 3 | Foxn1$^{+/+}$ | 0.84 | 0.58 | 1.10 | 0.13 | 41.00 | | | | 0.038 | 0.002 |
| Day 5 | Foxn1$^{+/+}$ | 0.25 | -0.04 | 0.54 | 0.14 | 41.00 | | | | | 0.946 |
| Day 7 | Foxn1$^{+/+}$ | 0.08 | -0.18 | 0.34 | 0.13 | 41.00 | | | | | |
| Day 0 | Foxn1$^{-/-}$ | 0.20 | -0.13 | 0.53 | 0.16 | 41.00 | 0.971 | 0.997 | 1.000 | 1.000 | 0.971 |
| Day 1 | Foxn1$^{-/-}$ | 0.04 | -0.22 | 0.30 | 0.13 | 41.00 | | 1.000 | 0.873 | 0.947 | 1.000 |
| Day 2 | Foxn1$^{-/-}$ | 0.10 | -0.19 | 0.39 | 0.14 | 41.00 | | | 0.976 | 0.995 | 1.000 |
| Day 3 | Foxn1$^{-/-}$ | 0.24 | -0.02 | 0.50 | 0.13 | 41.00 | | | | 1.000 | 0.873 |
| Day 5 | Foxn1$^{-/-}$ | 0.20 | -0.06 | 0.46 | 0.13 | 41.00 | | | | | 0.947 |
| Day 7 | Foxn1$^{-/-}$ | 0.04 | -0.22 | 0.30 | 0.13 | 41.00 | | | | | |

### MCP-1

**Supplementary Table 12. Comparisons by days for MCP-1**

| Day | group | lsmean | lower.CL | upper.CL | SE | df | Day 1 | Day 3 | Day 5 | Day 7 |
|---|---|---|---|---|---|---|---|---|---|---|
| Day 0 | BALB | 121.08 | -246.61 | 488.77 | 178.53 | 25.00 | 0.083 | 0.367 | 0.999 | 1.000 |
| Day 1 | BALB | 801.86 | 434.17 | 1169.55 | 178.53 | 25.00 | | 0.840 | 0.095 | 0.043 |
| Day 3 | BALB | 558.95 | 240.52 | 877.38 | 154.61 | 25.00 | | | 0.436 | 0.238 |
| Day 5 | BALB | 181.13 | -137.30 | 499.56 | 154.61 | 25.00 | | | | 0.994 |
| Day 7 | BALB | 92.84 | -225.59 | 411.27 | 154.61 | 25.00 | | | | |
| Day 0 | Nude | 210.32 | -157.37 | 578.01 | 178.53 | 25.00 | 0.804 | 0.148 | 0.980 | 1.000 |
| Day 1 | Nude | 521.10 | 70.77 | 971.42 | 218.65 | 25.00 | | 0.872 | 0.962 | 0.776 |
| Day 3 | Nude | 777.16 | 458.74 | 1095.59 | 154.61 | 25.00 | | | 0.300 | 0.104 |
| Day 5 | Nude | 342.39 | 23.96 | 660.82 | 154.61 | 25.00 | | | | 0.974 |
| Day 7 | Nude | 211.65 | -106.77 | 530.08 | 154.61 | 25.00 | | | | |

## CD68

**Supplementary Table 13. Comparisons by days for CD68**

| Day | group | lsmean | lower.CL | upper.CL | SE | df | Day 1 | Day 2 | Day 3 | Day 5 | Day 7 |
|---|---|---|---|---|---|---|---|---|---|---|---|
| Day 0 | Foxn1$^{+/+}$ | 1.63 | 0.17 | 3.10 | 0.73 | 44.00 | 1.000 | 0.985 | 1.000 | <0.001 | <0.001 |
| Day 1 | Foxn1$^{+/+}$ | 1.60 | 0.46 | 2.74 | 0.56 | 44.00 | | 0.977 | 0.999 | <0.001 | <0.001 |
| Day 2 | Foxn1$^{+/+}$ | 1.02 | -0.12 | 2.16 | 0.56 | 44.00 | | | 0.897 | <0.001 | <0.001 |
| Day 3 | Foxn1$^{+/+}$ | 1.86 | 0.72 | 3.00 | 0.56 | 44.00 | | | | <0.001 | <0.001 |
| Day 5 | Foxn1$^{+/+}$ | 10.94 | 9.80 | 12.08 | 0.56 | 44.00 | | | | | <0.001 |
| Day 7 | Foxn1$^{+/+}$ | 6.86 | 5.72 | 8.00 | 0.56 | 44.00 | | | | | |
| Day 0 | Foxn1$^{-/-}$ | 2.20 | 0.73 | 3.67 | 0.73 | 44.00 | 0.884 | 0.827 | 0.995 | <0.001 | <0.001 |
| Day 1 | Foxn1$^{-/-}$ | 1.20 | 0.06 | 2.34 | 0.56 | 44.00 | | 1.000 | 0.442 | <0.001 | <0.001 |
| Day 2 | Foxn1$^{-/-}$ | 1.08 | -0.06 | 2.22 | 0.56 | 44.00 | | | 0.355 | <0.001 | <0.001 |
| Day 3 | Foxn1$^{-/-}$ | 2.68 | 1.54 | 3.82 | 0.56 | 44.00 | | | | <0.001 | <0.001 |
| Day 5 | Foxn1$^{-/-}$ | 11.96 | 10.82 | 13.10 | 0.56 | 44.00 | | | | | <0.001 |
| Day 7 | Foxn1$^{-/-}$ | 7.16 | 6.02 | 8.30 | 0.56 | 44.00 | | | | | |

**Statistically significant differences for *Collagen I, Collagen III expression and* Hydroxyproline content in Foxn1$^{-/-}$ (CBy.Cg-Foxn1<nu>/cmdb) and Foxn1$^{+/+}$ (Balb/c/cmdb) mice during wound healing process**

### *Collagen I*

**Supplementary Table 14. Comparisons by group for *Collagen I***

| Day | group | lsmean | lower.CL | upper.CL | SE | Df | p.value |
|---|---|---|---|---|---|---|---|
| Day 0 | Foxn1$^{+/+}$ | 1.49 | -2.40 | 5.38 | 1.95 | 69.00 | <0.001 |
| Day 0 | Foxn1$^{-/-}$ | 14.28 | 10.39 | 18.17 | 1.95 | 69.00 | |
| Day 5 | Foxn1$^{+/+}$ | 22.24 | 18.35 | 26.13 | 1.95 | 69.00 | 0.003 |
| Day 5 | Foxn1$^{-/-}$ | 13.75 | 9.87 | 17.64 | 1.95 | 69.00 | |

**Supplementary Table 15. Comparisons by days for *Collagen I***

| Day | group | lsmean | lower.CL | upper.CL | SE | df | Day 1 | Day 3 | Day 5 | Day 7 | Day 14 | Day 21 |
|---|---|---|---|---|---|---|---|---|---|---|---|---|
| Day 0 | Foxn1$^{+/+}$ | 1.49 | -2.40 | 5.38 | 1.95 | 69.00 | 0.999 | 0.069 | <0.001 | <0.001 | 0.204 | 0.644 |
| Day 1 | Foxn1$^{+/+}$ | 2.85 | -1.04 | 6.73 | 1.95 | 69.00 | | 0.197 | <0.001 | <0.001 | 0.466 | 0.904 |
| Day 3 | Foxn1$^{+/+}$ | 9.90 | 5.64 | 14.16 | 2.13 | 69.00 | | | 0.001 | 0.054 | 0.997 | 0.836 |
| Day 5 | Foxn1$^{+/+}$ | 22.24 | 18.35 | 26.13 | 1.95 | 69.00 | | | | 0.840 | <0.001 | <0.001 |
| Day 7 | Foxn1$^{+/+}$ | 18.60 | 14.71 | 22.49 | 1.95 | 69.00 | | | | | 0.006 | <0.001 |
| Day 14 | Foxn1$^{+/+}$ | 8.17 | 4.29 | 12.06 | 1.95 | 69.00 | | | | | | 0.987 |
| Day 21 | Foxn1$^{+/+}$ | 6.06 | 2.18 | 9.95 | 1.95 | 69.00 | | | | | | |
| Day 0 | Foxn1$^{-/-}$ | 14.28 | 10.39 | 18.17 | 1.95 | 69.00 | 0.223 | 0.626 | 1.000 | 0.965 | 0.524 | 0.756 |
| Day 1 | Foxn1$^{-/-}$ | 7.72 | 3.84 | 11.61 | 1.95 | 69.00 | | 0.993 | 0.315 | 0.023 | 0.998 | 0.971 |
| Day 3 | Foxn1$^{-/-}$ | 9.63 | 5.74 | 13.52 | 1.95 | 69.00 | | | 0.746 | 0.134 | 1.000 | 1.000 |
| Day 5 | Foxn1$^{-/-}$ | 13.75 | 9.87 | 17.64 | 1.95 | 69.00 | | | | 0.917 | 0.649 | 0.855 |
| Day 7 | Foxn1$^{-/-}$ | 16.87 | 12.98 | 20.76 | 1.95 | 69.00 | | | | | 0.094 | 0.207 |
| Day 14 | Foxn1$^{-/-}$ | 9.20 | 5.31 | 13.09 | 1.95 | 69.00 | | | | | | 1.000 |
| Day 21 | Foxn1$^{-/-}$ | 10.21 | 6.32 | 14.09 | 1.95 | 69.00 | | | | | | |

### *Collagen III*

**Supplementary Table 16. Comparisons by group for *Collagen III***

| Day | group | lsmean | lower.CL | upper.CL | SE | df | p.value |
|---|---|---|---|---|---|---|---|
| Day 3 | Foxn1$^{+/+}$ | 3.24 | 2.07 | 4.41 | 0.58 | 67.00 | 0.001 |
| Day 3 | Foxn1$^{-/-}$ | 0.59 | -0.48 | 1.65 | 0.53 | 67.00 | |
| Day 5 | Foxn1$^{+/+}$ | 5.66 | 4.60 | 6.73 | 0.53 | 67.00 | <0.001 |
| Day 5 | Foxn1$^{-/-}$ | 0.65 | -0.42 | 1.71 | 0.53 | 67.00 | |
| Day 7 | Foxn1$^{+/+}$ | 2.87 | 1.80 | 3.93 | 0.53 | 67.00 | 0.001 |
| Day 7 | Foxn1$^{-/-}$ | 0.30 | -0.77 | 1.36 | 0.53 | 67.00 | |

**Supplementary Table 17. Comparisons by days for *Collagen III***

| Day | group | lsmean | lower.CL | upper.CL | SE | df | Day 1 | Day 3 | Day 5 | Day 7 | Day 14 | Day 21 |
|---|---|---|---|---|---|---|---|---|---|---|---|---|
| Day 0 | Foxn1$^{+/+}$ | 0.26 | -0.81 | 1.32 | 0.53 | 67.00 | 0.722 | 0.006 | <0.001 | 0.016 | 0.693 | 1.000 |
| Day 1 | Foxn1$^{+/+}$ | 1.42 | 0.35 | 2.48 | 0.53 | 67.00 | | 0.257 | <0.001 | 0.474 | 1.000 | 0.871 |
| Day 3 | Foxn1$^{+/+}$ | 3.24 | 2.07 | 4.41 | 0.58 | 67.00 | | | 0.047 | 0.999 | 0.279 | 0.014 |
| Day 5 | Foxn1$^{+/+}$ | 5.66 | 4.60 | 6.73 | 0.53 | 67.00 | | | | 0.007 | <0.001 | <0.001 |
| Day 7 | Foxn1$^{+/+}$ | 2.87 | 1.80 | 3.93 | 0.53 | 67.00 | | | | | 0.505 | 0.035 |
| Day 14 | Foxn1$^{+/+}$ | 1.45 | 0.39 | 2.52 | 0.53 | 67.00 | | | | | | 0.850 |
| Day 21 | Foxn1$^{+/+}$ | 0.47 | -0.59 | 1.54 | 0.53 | 67.00 | | | | | | |
| Day 0 | Foxn1$^{-/-}$ | 0.92 | -0.24 | 2.09 | 0.58 | 67.00 | 1.000 | 1.000 | 1.000 | 0.985 | 0.932 | 0.936 |
| Day 1 | Foxn1$^{-/-}$ | 1.06 | -0.01 | 2.12 | 0.53 | 67.00 | | 0.996 | 0.998 | 0.952 | 0.847 | 0.859 |
| Day 3 | Foxn1$^{-/-}$ | 0.59 | -0.48 | 1.65 | 0.53 | 67.00 | | | 1.000 | 1.000 | 0.993 | 0.993 |
| Day 5 | Foxn1$^{-/-}$ | 0.65 | -0.42 | 1.71 | 0.53 | 67.00 | | | | 0.999 | 0.988 | 0.988 |
| Day 7 | Foxn1$^{-/-}$ | 0.30 | -0.77 | 1.36 | 0.53 | 67.00 | | | | | 1.000 | 1.000 |
| Day 14 | Foxn1$^{-/-}$ | 0.07 | -0.99 | 1.14 | 0.53 | 67.00 | | | | | | 1.000 |

| Day | group | lsmean | lower.CL | upper.CL | SE | df | Day 1 | Day 3 | Day 5 | Day 7 | Day 14 | Day 21 |
|---|---|---|---|---|---|---|---|---|---|---|---|---|
| Day 21 | Foxn1$^{-/-}$ | 0.04 | -1.12 | 1.21 | 0.58 | 67.00 | | | | | | |

### Hydroxyproline content

**Supplementary Table 18. Comparisons by group for Hydroxyproline content**

| Day | group | lsmean | lower.CL | upper.CL | SE | df | p.value |
|---|---|---|---|---|---|---|---|
| Day 0 | Foxn1$^{+/+}$ | 4.25 | 3.29 | 5.21 | 0.48 | 41.00 | 0.002 |
| Day 0 | Foxn1$^{-/-}$ | 1.98 | 1.02 | 2.94 | 0.48 | 41.00 | |

### Statistically significant differences for *Tgfβ-1, Tgfβ-3, Mmp-9 and Timp-1 expression* in Foxn1$^{-/-}$ (CBy.Cg-Foxn1<nu>/cmdb) and Foxn1$^{+/+}$(Balb/c/cmdb) mice during wound healing process

#### *Tgfß-1*

**Supplementary Table 19. Comparisons by group for *Tgfß-1***

| Day | group | lsmean | lower.CL | upper.CL | SE | df | p.value |
|---|---|---|---|---|---|---|---|
| Day 5 | Foxn1$^{+/+}$ | 16.65 | 13.86 | 19.44 | 1.40 | 69.00 | <0.001 |
| Day 5 | Foxn1$^{-/-}$ | 8.32 | 5.53 | 11.11 | 1.40 | 69.00 | |

**Supplementary Table 20. Comparisons by days for *Tgfß-1***

| Day | group | lsmean | lower.CL | upper.CL | SE | df | Day 1 | Day 3 | Day 5 | Day 7 | Day 14 | Day 21 |
|---|---|---|---|---|---|---|---|---|---|---|---|---|
| Day 0 | Foxn1$^{+/+}$ | 12.71 | 9.92 | 15.50 | 1.40 | 69.00 | 1.000 | 0.901 | 0.430 | 0.998 | 0.118 | 0.744 |
| Day 1 | Foxn1$^{+/+}$ | 12.58 | 9.79 | 15.37 | 1.40 | 69.00 | | 0.876 | 0.389 | 0.996 | 0.137 | 0.783 |
| Day 3 | Foxn1$^{+/+}$ | 15.15 | 12.10 | 18.21 | 1.53 | 69.00 | | | 0.991 | 0.994 | 0.007 | 0.141 |
| Day 5 | Foxn1$^{+/+}$ | 16.65 | 13.86 | 19.44 | 1.40 | 69.00 | | | | 0.772 | <0.001 | 0.014 |
| Day 7 | Foxn1$^{+/+}$ | 13.78 | 10.99 | 16.57 | 1.40 | 69.00 | | | | | 0.030 | 0.401 |
| Day 14 | Foxn1$^{+/+}$ | 7.41 | 4.62 | 10.19 | 1.40 | 69.00 | | | | | | 0.897 |
| Day 21 | Foxn1$^{+/+}$ | 9.75 | 6.96 | 12.54 | 1.40 | 69.00 | | | | | | |
| Day 0 | Foxn1$^{-/-}$ | 11.67 | 8.88 | 14.46 | 1.40 | 69.00 | 0.512 | 0.901 | 0.623 | 1.000 | 0.694 | 0.475 |
| Day 1 | Foxn1$^{-/-}$ | 15.35 | 12.56 | 18.14 | 1.40 | 69.00 | | 0.993 | 0.012 | 0.615 | 0.016 | 0.006 |
| Day 3 | Foxn1$^{-/-}$ | 13.99 | 11.20 | 16.78 | 1.40 | 69.00 | | | 0.077 | 0.948 | 0.100 | 0.043 |
| Day 5 | Foxn1$^{-/-}$ | 8.32 | 5.53 | 11.11 | 1.40 | 69.00 | | | | 0.520 | 1.000 | 1.000 |
| Day 7 | Foxn1$^{-/-}$ | 11.98 | 9.19 | 14.77 | 1.40 | 69.00 | | | | | 0.593 | 0.378 |
| Day 14 | Foxn1$^{-/-}$ | 8.54 | 5.75 | 11.33 | 1.40 | 69.00 | | | | | | 1.000 |
| Day 21 | Foxn1$^{-/-}$ | 7.87 | 5.08 | 10.66 | 1.40 | 69.00 | | | | | | |

#### *Tgfß-3*

**Supplementary Table 21. Comparisons by group for *Tgfß-3***

| Day | group | lsmean | lower.CL | upper.CL | SE | df | p.value |
|---|---|---|---|---|---|---|---|
| Day 5 | Foxn1$^{+/+}$ | 1.98 | 0.55 | 3.42 | 0.72 | 69.00 | <0.001 |
| Day 5 | Foxn1$^{-/-}$ | 5.75 | 4.32 | 7.19 | 0.72 | 69.00 | |
| Day 7 | Foxn1$^{+/+}$ | 2.48 | 1.04 | 3.91 | 0.72 | 69.00 | 0.011 |
| Day 7 | Foxn1$^{-/-}$ | 5.12 | 3.69 | 6.56 | 0.72 | 69.00 | |
| Day 21 | Foxn1$^{+/+}$ | 1.63 | 0.19 | 3.07 | 0.72 | 69.00 | 0.021 |
| Day 21 | Foxn1$^{-/-}$ | 4.03 | 2.60 | 5.47 | 0.72 | 69.00 | |

**Supplementary Table 22. Comparisons by days for *Tgfß-3***

| Day | group | lsmean | lower.CL | upper.CL | SE | df | Day 1 | Day 3 | Day 5 | Day 7 | Day 14 | Day 21 |
|---|---|---|---|---|---|---|---|---|---|---|---|---|
| Day 0 | Foxn1$^{+/+}$ | 0.93 | -0.51 | 2.36 | 0.72 | 69.00 | 1.000 | 1.000 | 0.943 | 0.732 | 0.999 | 0.993 |
| Day 1 | Foxn1$^{+/+}$ | 0.94 | -0.50 | 2.38 | 0.72 | 69.00 | | 1.000 | 0.947 | 0.740 | 0.999 | 0.994 |
| Day 3 | Foxn1$^{+/+}$ | 1.11 | -0.46 | 2.69 | 0.79 | 69.00 | | | 0.983 | 0.861 | 1.000 | 0.999 |
| Day 5 | Foxn1$^{+/+}$ | 1.98 | 0.55 | 3.42 | 0.72 | 69.00 | | | | 0.999 | 0.997 | 1.000 |
| Day 7 | Foxn1$^{+/+}$ | 2.48 | 1.04 | 3.91 | 0.72 | 69.00 | | | | | 0.937 | 0.981 |
| Day 14 | Foxn1$^{+/+}$ | 1.40 | -0.04 | 2.83 | 0.72 | 69.00 | | | | | | 1.000 |
| Day 21 | Foxn1$^{+/+}$ | 1.63 | 0.19 | 3.07 | 0.72 | 69.00 | | | | | | |

| Day | group | lsmean | lower.CL | upper.CL | SE | df | Day 1 | Day 3 | Day 5 | Day 7 | Day 14 | Day 21 |
|---|---|---|---|---|---|---|---|---|---|---|---|---|
| Day 0 | Foxn1$^{-/-}$ | 2.22 | 0.78 | 3.66 | 0.72 | 69.00 | 0.997 | 1.000 | 0.015 | 0.080 | 1.000 | 0.566 |
| Day 1 | Foxn1$^{-/-}$ | 1.63 | 0.19 | 3.07 | 0.72 | 69.00 | | 0.991 | 0.002 | 0.017 | 0.955 | 0.232 |
| Day 3 | Foxn1$^{-/-}$ | 2.36 | 0.92 | 3.79 | 0.72 | 69.00 | | | 0.022 | 0.110 | 1.000 | 0.653 |
| Day 5 | Foxn1$^{-/-}$ | 5.75 | 4.32 | 7.19 | 0.72 | 69.00 | | | | 0.996 | 0.047 | 0.625 |
| Day 7 | Foxn1$^{-/-}$ | 5.12 | 3.69 | 6.56 | 0.72 | 69.00 | | | | | 0.198 | 0.935 |
| Day 14 | Foxn1$^{-/-}$ | 2.64 | 1.20 | 4.07 | 0.72 | 69.00 | | | | | | 0.815 |
| Day 21 | Foxn1$^{-/-}$ | 4.03 | 2.60 | 5.47 | 0.72 | 69.00 | | | | | | |

### *Mmp-9*

**Supplementary Table 23. Comparisons by group for *Mmp-9***

| Day | group | lsmean | lower.CL | upper.CL | SE | df | p.value |
|---|---|---|---|---|---|---|---|
| Day 5 | Foxn1$^{+/+}$ | 39.86 | 30.49 | 49.22 | 4.70 | 69.00 | <0.001 |
| Day 5 | Foxn1$^{-/-}$ | 12.49 | 3.12 | 21.85 | 4.70 | 69.00 | |

**Supplementary Table 24. Comparisons by days for *Mmp-9***

| Day | group | lsmean | lower.CL | upper.CL | SE | df | Day 1 | Day 3 | Day 5 | Day 7 | Day 14 | Day 21 |
|---|---|---|---|---|---|---|---|---|---|---|---|---|
| Day 0 | Foxn1$^{+/+}$ | 7.01 | -2.36 | 16.37 | 4.70 | 69.00 | 0.193 | 0.624 | <0.001 | 0.996 | 0.998 | 0.998 |
| Day 1 | Foxn1$^{+/+}$ | 23.30 | 13.93 | 32.66 | 4.70 | 69.00 | | 0.995 | 0.178 | 0.522 | 0.053 | 0.056 |
| Day 3 | Foxn1$^{+/+}$ | 18.79 | 8.52 | 29.05 | 5.14 | 69.00 | | | 0.051 | 0.922 | 0.295 | 0.307 |
| Day 5 | Foxn1$^{+/+}$ | 39.86 | 30.49 | 49.22 | 4.70 | 69.00 | | | | <0.001 | <0.001 | <0.001 |
| Day 7 | Foxn1$^{+/+}$ | 11.04 | 1.67 | 20.41 | 4.70 | 69.00 | | | | | 0.903 | 0.912 |
| Day 14 | Foxn1$^{+/+}$ | 3.28 | -6.09 | 12.64 | 4.70 | 69.00 | | | | | | 1.000 |
| Day 21 | Foxn1$^{+/+}$ | 3.44 | -5.93 | 12.80 | 4.70 | 69.00 | | | | | | |
| Day 0 | Foxn1$^{-/-}$ | 17.11 | 7.74 | 26.48 | 4.70 | 69.00 | 0.297 | 1.000 | 0.992 | 0.965 | 0.873 | 0.904 |
| Day 1 | Foxn1$^{-/-}$ | 31.87 | 22.50 | 41.24 | 4.70 | 69.00 | | 0.241 | 0.067 | 0.036 | 0.015 | 0.019 |
| Day 3 | Foxn1$^{-/-}$ | 16.34 | 6.98 | 25.71 | 4.70 | 69.00 | | | 0.997 | 0.982 | 0.916 | 0.940 |
| Day 5 | Foxn1$^{-/-}$ | 12.49 | 3.12 | 21.85 | 4.70 | 69.00 | | | | 1.000 | 0.998 | 0.999 |
| Day 7 | Foxn1$^{-/-}$ | 10.88 | 1.51 | 20.24 | 4.70 | 69.00 | | | | | 1.000 | 1.000 |
| Day 14 | Foxn1$^{-/-}$ | 8.83 | -0.54 | 18.20 | 4.70 | 69.00 | | | | | | 1.000 |
| Day 21 | Foxn1$^{-/-}$ | 9.36 | -0.00 | 18.73 | 4.70 | 69.00 | | | | | | |

### *Timp-1*

**Supplementary Table 25. Comparisons by group for *Timp-1***

| Day | group | lsmean | lower.CL | upper.CL | SE | df | p.value |
|---|---|---|---|---|---|---|---|
| Day 1 | Foxn1$^{+/+}$ | 70.52 | 55.22 | 85.81 | 7.66 | 67.00 | <0.001 |
| Day 1 | Foxn1$^{-/-}$ | 30.23 | 14.93 | 45.53 | 7.66 | 67.00 | |
| Day 3 | Foxn1$^{+/+}$ | 54.32 | 37.56 | 71.07 | 8.39 | 67.00 | 0.002 |
| Day 3 | Foxn1$^{-/-}$ | 17.79 | 2.49 | 33.08 | 7.66 | 67.00 | |
| Day 5 | Foxn1$^{+/+}$ | 39.08 | 23.78 | 54.37 | 7.66 | 67.00 | 0.003 |
| Day 5 | Foxn1$^{-/-}$ | 6.21 | -9.08 | 21.51 | 7.66 | 67.00 | |

**Supplementary Table 26. Comparisons by days for *Timp-1***

| Day | group | lsmean | lower.CL | upper.CL | SE | df | Day 1 | Day 3 | Day 5 | Day 7 | Day 14 | Day 21 |
|---|---|---|---|---|---|---|---|---|---|---|---|---|
| Day 0 | Foxn1$^{+/+}$ | 20.66 | 5.36 | 35.96 | 7.66 | 67.00 | <0.001 | 0.061 | 0.619 | 0.963 | 0.623 | 0.526 |
| Day 1 | Foxn1$^{+/+}$ | 70.52 | 55.22 | 85.81 | 7.66 | 67.00 | | 0.786 | 0.071 | <0.001 | <0.001 | <0.001 |
| Day 3 | Foxn1$^{+/+}$ | 54.32 | 37.56 | 71.07 | 8.39 | 67.00 | | | 0.830 | 0.004 | <0.001 | <0.001 |
| Day 5 | Foxn1$^{+/+}$ | 39.08 | 23.78 | 54.37 | 7.66 | 67.00 | | | | 0.128 | 0.019 | 0.012 |
| Day 7 | Foxn1$^{+/+}$ | 10.36 | -4.93 | 25.66 | 7.66 | 67.00 | | | | | 0.989 | 0.973 |
| Day 14 | Foxn1$^{+/+}$ | 2.32 | -12.98 | 17.61 | 7.66 | 67.00 | | | | | | 1.000 |
| Day 21 | Foxn1$^{+/+}$ | 0.71 | -14.59 | 16.00 | 7.66 | 67.00 | | | | | | |
| Day 0 | Foxn1$^{-/-}$ | 3.50 | -13.26 | 20.25 | 8.39 | 67.00 | 0.236 | 0.869 | 1.000 | 1.000 | 1.000 | 1.000 |
| Day 1 | Foxn1$^{-/-}$ | 30.23 | 14.93 | 45.53 | 7.66 | 67.00 | | 0.911 | 0.301 | 0.174 | 0.135 | 0.172 |
| Day 3 | Foxn1$^{-/-}$ | 17.79 | 2.49 | 33.08 | 7.66 | 67.00 | | | 0.935 | 0.821 | 0.757 | 0.791 |

| Day | group | lsmean | lower.CL | upper.CL | SE | df | Day 1 | Day 3 | Day 5 | Day 7 | Day 14 | Day 21 |
|---|---|---|---|---|---|---|---|---|---|---|---|---|
| Day 5 | Foxn1$^{-/-}$ | 6.21 | -9.08 | 21.51 | 7.66 | 67.00 | | | | 1.000 | 1.000 | 1.000 |
| Day 7 | Foxn1$^{-/-}$ | 3.07 | -12.23 | 18.36 | 7.66 | 67.00 | | | | | 1.000 | 1.000 |
| Day 14 | Foxn1$^{-/-}$ | 1.78 | -13.51 | 17.08 | 7.66 | 67.00 | | | | | | 1.000 |
| Day 21 | Foxn1$^{-/-}$ | 1.69 | -15.06 | 18.45 | 8.39 | 67.00 | | | | | | |

### Statistically significant differences for *α-Sma, and Collagen III mRNA expression in* control, Tgfβ-1 or Tgfβ-3 treated dermal fibroblasts

#### *α-Sma*

**Supplementary Table 27. Comparisons by conditions for *α-Sma***

| condition | group | lsmean | lower.CL | upper.CL | SE | df | Tgfß-1 treated | Tgfß-3 treated |
|---|---|---|---|---|---|---|---|---|
| ctrl | Foxn1$^{+/+}$ | 8.49 | 7.95 | 9.03 | 0.26 | 24.00 | <0.001 | <0.001 |
| Tgfß-1 treated | Foxn1$^{+/+}$ | 10.38 | 9.83 | 10.92 | 0.26 | 24.00 | | 0.055 |
| Tgfß-3 treated | Foxn1$^{+/+}$ | 11.29 | 10.75 | 11.84 | 0.26 | 24.00 | | |
| ctrl | Foxn1$^{-/-}$ | 8.58 | 8.03 | 9.12 | 0.26 | 24.00 | <0.001 | <0.001 |
| Tgfß-1 treated | Foxn1$^{-/-}$ | 10.26 | 9.72 | 10.80 | 0.26 | 24.00 | | 0.070 |
| Tgfß-3 treated | Foxn1$^{-/-}$ | 11.13 | 10.59 | 11.68 | 0.26 | 24.00 | | |

#### *Collagen III*

**Supplementary Table 28. Comparisons by group for *Collagen III***

| Day | group | lsmean | lower.CL | upper.CL | SE | df | p.value |
|---|---|---|---|---|---|---|---|
| Day 3 | Foxn1$^{+/+}$ | 3.24 | 2.07 | 4.41 | 0.58 | 67.00 | 0.001 |
| Day 3 | Foxn1$^{-/-}$ | 0.59 | -0.48 | 1.65 | 0.53 | 67.00 | |
| Day 5 | Foxn1$^{+/+}$ | 5.66 | 4.60 | 6.73 | 0.53 | 67.00 | <0.001 |
| Day 5 | Foxn1$^{-/-}$ | 0.65 | -0.42 | 1.71 | 0.53 | 67.00 | |
| Day 7 | Foxn1$^{+/+}$ | 2.87 | 1.80 | 3.93 | 0.53 | 67.00 | 0.001 |
| Day 7 | Foxn1$^{-/-}$ | 0.30 | -0.77 | 1.36 | 0.53 | 67.00 | |

**Supplementary Table 29. Comparisons by days for *Collagen III***

| Day | group | lsmean | lower.CL | upper.CL | SE | df | Day 1 | Day 3 | Day 5 | Day 7 | Day 14 | Day 21 |
|---|---|---|---|---|---|---|---|---|---|---|---|---|
| Day 0 | Foxn1$^{+/+}$ | 0.26 | -0.81 | 1.32 | 0.53 | 67.00 | 0.722 | 0.006 | <0.001 | 0.016 | 0.693 | 1.000 |
| Day 1 | Foxn1$^{+/+}$ | 1.42 | 0.35 | 2.48 | 0.53 | 67.00 | | 0.257 | <0.001 | 0.474 | 1.000 | 0.871 |
| Day 3 | Foxn1$^{+/+}$ | 3.24 | 2.07 | 4.41 | 0.58 | 67.00 | | | 0.047 | 0.999 | 0.279 | 0.014 |
| Day 5 | Foxn1$^{+/+}$ | 5.66 | 4.60 | 6.73 | 0.53 | 67.00 | | | | 0.007 | <0.001 | <0.001 |
| Day 7 | Foxn1$^{+/+}$ | 2.87 | 1.80 | 3.93 | 0.53 | 67.00 | | | | | 0.505 | 0.035 |
| Day 14 | Foxn1$^{+/+}$ | 1.45 | 0.39 | 2.52 | 0.53 | 67.00 | | | | | | 0.850 |
| Day 21 | Foxn1$^{+/+}$ | 0.47 | -0.59 | 1.54 | 0.53 | 67.00 | | | | | | |
| Day 0 | Foxn1$^{-/-}$ | 0.92 | -0.24 | 2.09 | 0.58 | 67.00 | 1.000 | 1.000 | 1.000 | 0.985 | 0.932 | 0.936 |
| Day 1 | Foxn1$^{-/-}$ | 1.06 | -0.01 | 2.12 | 0.53 | 67.00 | | 0.996 | 0.998 | 0.952 | 0.847 | 0.859 |
| Day 3 | Foxn1$^{-/-}$ | 0.59 | -0.48 | 1.65 | 0.53 | 67.00 | | | 1.000 | 1.000 | 0.993 | 0.993 |
| Day 5 | Foxn1$^{-/-}$ | 0.65 | -0.42 | 1.71 | 0.53 | 67.00 | | | | 0.999 | 0.988 | 0.988 |
| Day 7 | Foxn1$^{-/-}$ | 0.30 | -0.77 | 1.36 | 0.53 | 67.00 | | | | | 1.000 | 1.000 |
| Day 14 | Foxn1$^{-/-}$ | 0.07 | -0.99 | 1.14 | 0.53 | 67.00 | | | | | | 1.000 |
| Day 21 | Foxn1$^{-/-}$ | 0.04 | -1.12 | 1.21 | 0.58 | 67.00 | | | | | | |

### Statistically significant differences for collagen I protein content analyzed in the media collected from cultured Foxn1$^{-/-}$ or Foxn1$^{+/+}$ dermal fibroblasts (DFs) treated with TGFβ-1 or TGFβ-3 or non-treated (control).

#### Collagen I

**Supplementary Table 30. Comparisons by group for Collagen I Elisa Test**

| condition | time | group | lsmean | lower.CL | upper.CL | SE | df | p.value |
|---|---|---|---|---|---|---|---|---|
| tgfß-1 | 72H | Foxn1$^{+/+}$ | 6.90 | 5.65 | 8.15 | 0.62 | 46.00 | 0.016 |
| tgfß-1 | 72H | Foxn1$^{-/-}$ | 9.09 | 7.84 | 10.34 | 0.62 | 46.00 | |
| tgfß-3 | 48H | Foxn1$^{+/+}$ | 4.49 | 3.24 | 5.74 | 0.62 | 46.00 | <0.001 |

| condition | time | group | lsmean | lower.CL | upper.CL | SE | df | p.value |
|---|---|---|---|---|---|---|---|---|
| tgfß-3 | 48H | $Foxn1^{-/-}$ | 8.52 | 7.27 | 9.77 | 0.62 | 46.00 | |
| tgfß-3 | 72H | $Foxn1^{+/+}$ | 7.84 | 6.59 | 9.09 | 0.62 | 46.00 | <0.001 |
| tgfß-3 | 72H | $Foxn1^{-/-}$ | 11.95 | 10.70 | 13.20 | 0.62 | 46.00 | |

**Supplementary Table 31. Comparisons by conditions for Collagen I Elisa Test**

| condition | time | group | lsmean | lower.CL | upper.CL | SE | df | tgfß-1 | tgfß-3 |
|---|---|---|---|---|---|---|---|---|---|
| control | 48H | $Foxn1^{+/+}$ | 7.67 | 6.22 | 9.11 | 0.72 | 46.00 | 0.706 | 0.005 |
| tgfß-1 | 48H | $Foxn1^{+/+}$ | 8.42 | 7.17 | 9.67 | 0.62 | 46.00 | | <0.001 |
| control | 72H | $Foxn1^{+/+}$ | 9.42 | 7.98 | 10.86 | 0.72 | 46.00 | 0.028 | 0.230 |
| tgfß-1 | 72H | $Foxn1^{-/-}$ | 9.09 | 7.84 | 10.34 | 0.62 | 46.00 | | 0.006 |

**Supplementary Table 32. Comparisons by time for Collagen I Elisa Test**

| condition | time | group | lsmean | lower.CL | upper.CL | SE | df | 48H | 72H |
|---|---|---|---|---|---|---|---|---|---|
| control | 24H | $Foxn1^{+/+}$ | 2.75 | 1.31 | 4.19 | 0.72 | 46.00 | <0.001 | <0.001 |
| tgfß-1 | 24H | $Foxn1^{+/+}$ | 2.82 | 1.38 | 4.26 | 0.72 | 46.00 | <0.001 | <0.001 |
| tgfß-3 | 24H | $Foxn1^{+/+}$ | 3.60 | 2.35 | 4.85 | 0.62 | 46.00 | 0.573 | <0.001 |
| tgfß-3 | 48H | $Foxn1^{+/+}$ | 4.49 | 3.24 | 5.74 | 0.62 | 46.00 | | 0.001 |
| control | 24H | $Foxn1^{-/-}$ | 3.37 | 1.93 | 4.81 | 0.72 | 46.00 | 0.003 | <0.001 |
| control | 48H | $Foxn1^{-/-}$ | 6.96 | 5.52 | 8.40 | 0.72 | 46.00 | | <0.001 |
| tgfß-1 | 24H | $Foxn1^{-/-}$ | 3.73 | 2.29 | 5.17 | 0.72 | 46.00 | <0.001 | <0.001 |
| tgfß-3 | 24H | $Foxn1^{-/-}$ | 3.70 | 2.45 | 4.95 | 0.62 | 46.00 | <0.001 | <0.001 |
| tgfß-3 | 48H | $Foxn1^{-/-}$ | 8.52 | 7.27 | 9.77 | 0.62 | 46.00 | | <0.001 |

### Statistically significant differences for *Foxn1, Hif-1α* and *Fih-1 (Hif1-an)* mRNA expression collected from $Foxn1^{-/-}$ and $Foxn1^{+/+}$ keratinocytes cultured: under different oxygen availability (21%$O_2$ or 1%$O_2$), time of culture (12h vs 24h) and method (transduced with Ad-Foxn1 or Ad-GFP or non-transduced).

#### *Foxn1*

**Supplementary Table 33. Comparisons by time 12h vs 24h of culture – $Foxn1^{+/+}$ keratinocytes.**

| $O_2$ | time | lsmean | lower.CL | upper.CL | SE | df | p.value |
|---|---|---|---|---|---|---|---|
| 21% | 12 h | 0.03 | 0.01 | 0.06 | 0.01 | 11.00 | 0.001 |
| 21% | 24 h | 0.11 | 0.08 | 0.13 | 0.01 | 11.00 | |

**Supplementary Table 34. Comparisons by $O_2$ levels 1%$O_2$ vs 21%$O_2$ – $Foxn1^{+/+}$ keratinocytes.**

| $O_2$ | time | lsmean | lower.CL | upper.CL | SE | df | p.value |
|---|---|---|---|---|---|---|---|
| 1% | 12 h | 0.08 | 0.06 | 0.11 | 0.01 | 11.00 | 0.013 |
| 21% | 12 h | 0.03 | 0.01 | 0.06 | 0.01 | 11.00 | |

**Supplementary Table 35. Comparisons by method: Ad-Foxn1 or Ad-GFP transfected – $Foxn1^{-/-}$ keratinocytes.**

| $O_2$ | method | lsmean | lower.CL | upper.CL | SE | df | p.value |
|---|---|---|---|---|---|---|---|
| 1% | Ad-Foxn1 | 71.81 | 67.81 | 75.81 | 1.74 | 8.00 | <0.001 |
| 1% | Ad-GFP | 0.02 | -3.98 | 4.03 | 1.74 | 8.00 | |
| 21% | Ad-Foxn1 | 32.55 | 28.55 | 36.56 | 1.74 | 8.00 | <0.001 |
| 21% | Ad-GFP | 0.02 | -3.98 | 4.02 | 1.74 | 8.00 | |

**Supplementary Table 36. Comparisons by $O_2$ levels 1%$O_2$ vs 21%$O_2$ – $Foxn1^{-/-}$ keratinocytes.**

| $O_2$ | method | lsmean | lower.CL | upper.CL | SE | df | p.value |
|---|---|---|---|---|---|---|---|
| 1% | Ad-Foxn1 | 71.81 | 67.81 | 75.81 | 1.74 | 8.00 | <0.001 |
| 21% | Ad-Foxn1 | 32.55 | 28.55 | 36.56 | 1.74 | 8.00 | |

### *Hif-1α*

**Supplementary Table 37. Comparisons by time for *Hif-1α* - Foxn1$^{+/+}$ keratinocytes**

| $O_2$ | time | lsmean | lower.CL | upper.CL | SE | df | p.value |
|---|---|---|---|---|---|---|---|
| 1% | 12 h | 4.11 | 3.28 | 4.94 | 0.38 | 11.00 | <0.001 |
| 1% | 24 h | 6.82 | 5.86 | 7.77 | 0.43 | 11.00 | |
| 21% | 12 h | 5.36 | 4.53 | 6.19 | 0.38 | 11.00 | <0.001 |
| 21% | 24 h | 8.79 | 7.96 | 9.62 | 0.38 | 11.00 | |

**Supplementary Table 38. Comparisons by $O_2$ level for *Hif-1α* - Foxn1$^{+/+}$ keratinocytes**

| $O_2$ | time | lsmean | lower.CL | upper.CL | SE | df | p.value |
|---|---|---|---|---|---|---|---|
| 1% | 12 h | 4.11 | 3.28 | 4.94 | 0.38 | 11.00 | 0.039 |
| 21% | 12 h | 5.36 | 4.53 | 6.19 | 0.38 | 11.00 | |
| 1% | 24 h | 6.82 | 5.86 | 7.77 | 0.43 | 11.00 | 0.006 |
| 21% | 24 h | 8.79 | 7.96 | 9.62 | 0.38 | 11.00 | |

**Supplementary Table 39. Comparisons by method for *Hif-1α* – Foxn1$^{-/-}$ keratinocytes**

| $O_2$ | method | lsmean | lower.CL | upper.CL | SE | df | p.value |
|---|---|---|---|---|---|---|---|
| 21% | Ad-Foxn1 | 6.50 | 5.13 | 7.86 | 0.59 | 8.00 | <0.001 |
| 21% | Ad-GFP | 11.26 | 9.90 | 12.62 | 0.59 | 8.00 | |

**Supplementary Table 40. Comparisons by $O_2$ level for *Hif-1α* – Foxn1$^{-/-}$ keratinocytes**

| $O_2$ | method | lsmean | lower.CL | upper.CL | SE | df | p.value |
|---|---|---|---|---|---|---|---|
| 1% | Ad-Foxn1 | 8.75 | 7.39 | 10.12 | 0.59 | 8.00 | 0.027 |
| 21% | Ad-Foxn1 | 6.50 | 5.13 | 7.86 | 0.59 | 8.00 | |
| 1% | Ad-GFP | 8.22 | 6.86 | 9.58 | 0.59 | 8.00 | 0.007 |
| 21% | Ad-GFP | 11.26 | 9.90 | 12.62 | 0.59 | 8.00 | |

**Supplementary Table 41. Comparisons by method for *Hif-1α* - after 24 h**

| $O_2$ | method | lsmean | lower.CL | upper.CL | SE | df | Ad-GFP | B6 mice |
|---|---|---|---|---|---|---|---|---|
| 21% | Ad-Foxn1 | 6.50 | 5.18 | 7.81 | 0.61 | 13.00 | <0.001 | 0.034 |
| 21% | Ad-GFP | 11.26 | 9.95 | 12.58 | 0.61 | 13.00 | | 0.023 |

**Supplementary Table 42. Comparisons by $O_2$ level for *Hif-1α* - after 24 h**

| $O_2$ | method | lsmean | lower.CL | upper.CL | SE | df | p.value |
|---|---|---|---|---|---|---|---|
| 1% | Ad-Foxn1 | 8.75 | 7.44 | 10.07 | 0.61 | 13.00 | 0.021 |
| 21% | Ad-Foxn1 | 6.50 | 5.18 | 7.81 | 0.61 | 13.00 | |
| 1% | Ad-GFP | 8.22 | 6.90 | 9.53 | 0.61 | 13.00 | 0.004 |
| 21% | Ad-GFP | 11.26 | 9.95 | 12.58 | 0.61 | 13.00 | |
| 1% | Foxn1$^{+/+}$ | 6.82 | 5.50 | 8.13 | 0.61 | 13.00 | 0.029 |
| 21% | Foxn1$^{+/+}$ | 8.79 | 7.65 | 9.93 | 0.53 | 13.00 | |

### *Fih-1 (Hif-1an)*

**Supplementary Table 43. Comparisons by method for *Fih-1* (*Hif1-an*) – Foxn1$^{-/-}$ keratinocytes**

| $O_2$ | method | lsmean | lower.CL | upper.CL | SE | df | p.value |
|---|---|---|---|---|---|---|---|
| 1% | Ad-Foxn1 | 8.61 | 8.14 | 9.08 | 0.20 | 8.00 | <0.001 |
| 1% | Ad-GFP | 10.45 | 9.98 | 10.92 | 0.20 | 8.00 | |

**Supplementary Table 44. Comparisons by $O_2$ level for *Fih-1* (*Hif1-an*) – Foxn1$^{-/-}$ keratinocytes**

| $O_2$ | method | lsmean | lower.CL | upper.CL | SE | df | p.value |
|---|---|---|---|---|---|---|---|
| 1% | Ad-Foxn1 | 8.61 | 8.14 | 9.08 | 0.20 | 8.00 | 0.002 |
| 21% | Ad-Foxn1 | 9.95 | 9.48 | 10.42 | 0.20 | 8.00 | |
